# Mutant CHCHD10 disrupts cytochrome *c* oxidation and activates mitochondrial retrograde signaling in a model of cardiomyopathy

**DOI:** 10.1101/2025.08.18.668848

**Authors:** Márcio Augusto Campos-Ribeiro, Erminia Donnarumma, Hendrik Nolte, Paul Cobine, Elodie Vimont, Dusanka Milenkovic, Juan Diego Hernandez-Camacho, Francina Langa Vives, Etienne Kornobis, Esthel Pénard, Thomas Langer, Véronique Paquis-Flucklinger, Timothy Wai

## Abstract

Mutations in *CHCHD10*, a mitochondrial intermembrane space (IMS) protein implicated in proteostasis and cristae maintenance, cause multi-systemic mitochondrial disease. Heterozygous *Chchd10* knock-in mice modeling the human *CHCHD10^S59L^* variant associated with Amyotrophic Lateral Sclerosis and Frontotemporal Dementia (ALS-FTD) develop a mitochondrial cardiomyopathy driven by CHCHD10 insolubility and aggregation, which is associated with chronic activation of the mitochondrial integrated stress response (mtISR). Here, we demonstrate that cardiac dysfunction in *Chchd10^S55L/+^* mice carrying the orthologous pathogenic variant is associated with dual defects originating at the onset of disease: (1) early bioenergetic dysfunction linked to defects in the mitochondrial copper homeostasis and the oxidation of cytochrome *c* and (2) maladaptive mtISR signaling via the OMA1-DELE1-HRI axis. Using *Oma1^E324Q/E324Q^*knock-in mice, we show that the catalytic inactivation of the mitochondrial protease OMA1 in *Chchd10^S55L/+^* mice delays cardiomyopathy onset without rescuing CHCHD10 insolubility, proteomic remodeling, cristae defects or OXPHOS impairment, demonstrating that mtISR can be uncoupled from the bioenergetic collapse triggered by mutant CHCHD10. Proteomic profiling of soluble and insoluble mitochondrial proteins in *Chchd10^S55L/+^* mice reveals wide-spread disruptions of mitochondrial proteostasis, including IMS proteins involved in cytochrome *c* biogenesis. Defective respiration in mutant mitochondria could be rescued by the exogenous addition of cytochrome *c*, pinpointing IMS proteostasis disruption as a key pathogenic mechanism. Our work reveals that mutant CHCHD10 insolubility compromises metabolic resilience by impairing both mitochondrial bioenergetics and stress adaptation, offering new perspectives for the development of therapeutic targets.

## Introduction

Mitochondria are multi-functional organelles that execute essential biosynthetic and signaling functions that govern the life and death of the cell^1^. Inherited defects in mitochondria cause Mitochondrial Diseases (MD), which are rare, clinically heterogeneous and multi-systemic disorders caused by pathogenic variants in genes that code for ∼ 400 out of the >1200 mitochondrial proteins that have been identified to date^2,3^. The tissue-specific nature of MD has long been proposed to reflect the differential bioenergetic and metabolic requirements and tissue-specific biochemical thresholds^4^. More recent studies exploring the functional impact of downstream stress signaling triggered by mitochondrial dysfunction have highlighted the physiological relevance of cell signaling for disease etiology, trajectory and severity^5,6^. The mitochondrial integrated stress response (mtISR)^7^ depends on the proteolytic cleavage and maturation of DAP3 Binding Cell Death Enhancer 1 (DELE1) within mitochondria by the stress-induced mitochondrial protease OMA1^8,9^, which enables translocation of DELE1 to the cytosol where it activates the heme-regulated inhibitor kinase HRI, leading to cytosolic translational attenuation and selective upregulation of stress-responsive genes encoding chaperones, transport proteins, and proteases by <u>A</u>ctivating <u>T</u>ranscription <u>F</u>actors like ATF4^10^.

CHCHD10 is a small 14kDa protein that belongs to the family of CHCHD proteins, which are characterized by the presence of a CHCH domain, which consists of two helix-coil-helix motifs connected by a loop. The twin Cx(9)C motif (C-X₉-C-X₂-C-X₉-C) contains four conserved cysteines that are required for intramolecular disulfide bridges that are critical for protein stability and function. CHCHD proteins are translated on cytosolic ribosomes and imported into the intermembrane space (IMS) via the disulfide relay system catalyzed by the copper– and zinc-dependent protein (encoded by *CHCHD4*) and the Mitochondrial FAD-linked sulfhydryl oxidase ERV1 (encoded by *GFER*)^11^. MIA40 forms a redox cycle with cysteine motif-containing client proteins destined for IMS import, which are initially translocated in a reduced state. Upon entry, the oxidized form of MIA40 forms a transient intermolecular disulfide intermediate with the reduced precursor, facilitating the transfer of disulfide bonds. Electrons passed from MIA40 to ERV1, which reduces FAD to FADH_2_, are then passed on to cytochrome *c* or molecular oxygen (O_2_) to complete the redox cycle. Defects in the MIA40/ERV1 import pathway can impinge on the biogenesis of client proteins, which play roles in various intramitochondrial processes including the assembly of enzymes of the electron transport chain (ETC), maintenance of the MICOS complex and cristae morphology, and mitochondrial proteostasis^11^. Inborn errors in ERV1/GFER are associated with multi-systemic MD characterized by combined respiratory chain deficiency and disordered cristae^12,13^. Pathogenic variants in the mitochondrial protein coiled-coil-helix-coiled-coil-helix domain-containing protein 10 (CHCHD10) were initially linked to autosomal dominant (AD) amyotrophic lateral sclerosis and frontotemporal dementia (ALS-FTD)^14–16^ and later studies identified mutations in patients suffering from Charcot-Marie-Tooth neuropathy^17^, spinal muscular atrophy^18^, mitochondrial myopathy, and cardiomyopathy^19^. Transgenic mouse models for these pathogenic variants recapitulate some of the cellular dysfunctions described in patients, yet there are conflicting hypothesis regarding the precise molecular defects underscoring disease onset^20–23^.

Toxic gain-of-function mechanisms underlying the expressivity of the AD *CHCHD10* variant S59L (OMIM: #615911) promotes CHCHD10 insolubility, aggregation, and accumulation, consistent with atomic-level structural modeling studies suggesting a greater propensity for oligomerization^24^. For CHCHD10^S59L^, insolubility occurs within the IMS and promotes the accumulation and insolubility of its interaction partner CHCHD2 in both patients-derived cells and mouse models^19,25–28^. In *CHCHD10^S59L/+^* patients, muscle biopsies showed ragged-red fibers, cytochrome *c* oxidase (COX)-negative fibers, and mitochondrial DNA (mtDNA) deletions that have been proposed to result from mtISR-dependent nucleotide imbalance^21^. Heterozygous *Chchd10^S55L/+^* knock-in mice (carrying the S59L murine equivalent) develop a tissue-specific MD characterized by an early onset cardiomyopathy and inability to gain weight, followed by a late-onset neuromuscular decline and death around 1 year of age. In these animals, CHCHD10 accumulates and aggregates in affected tissues, leading to mtISR induction and culminating in extensive proteomic, morphological and ultrastructural remodeling of mitochondria^21,25,26,29^. The prevailing model proposes that mtISR induction triggers progressive metabolic and late-stage OXPHOS decline in *Chchd10^S55L/+^*knock-in mice through the remodeling of pathways governed by proteins encoded by ISR target genes^21^. Efforts to blunt the mtISR through the whole-body *Oma1* deletion in *Chchd10^G54R/+^*knock-in mice, which model an AD mitochondrial myopathy caused by the G58R gain-of-function variant, revealed OMA1 and mutant CHCHD10 to be synthetically lethal^19^. In this model, tissue-specific ablation of *Oma1* and/or *Dele1* exacerbated myopathy, leading to the opposite conclusion that the mtISR is required for tissue homeostasis^22^, at least in skeletal muscle. Hence, the functional impact of the mtISR remains controversial, with its inhibition in models of mitochondrial dysfunction beyond CHCHD10 reported to be both protective and maladaptive in vivo^30–35^.

In this study, we set out to characterize the relationship between CHCHD10 protein insolubility caused by the S59L mutation, the mtISR signaling pathway, and mitochondrial bioenergetics in the heart. We confirmed that CHCHD10 protein insolubility and mtISR induction precede cardiac dysfunction and are therefore a candidate trigger for disease onset in *CHCHD10^S55L/+^* mice. To determine the relevance of the mtISR, we introduced the catalytically-inactivating mutation E324Q in the *Oma1* gene of *CHCHD10^S55L/+^*mice, which blunted mtISR signaling and delayed cardiomyopathy without rescuing early-onset defects in CHCHD10 insolubility, cristae structure, cytochrome *c* oxidation, nor mtDNA depletion. Biochemical and OMICs-based studies of cardiac mitochondria from *CHCHD10^S55L/+^*mice revealed defects in cytochrome *c* oxidation capacity, mitochondrial copper levels, and mitochondrial respiration that manifest at or before disease onset, which is in contrast with previous studies proposing OXPHOS defect to be a late-stage consequence of ISR-dependent rewiring of mitochondria^21,28^. Importantly, in vitro supplementation of exogenous cytochrome *c* was able to rescue impaired respiration in mitoplasts from *CHCHD10^S55L/+^* hearts, which were found to be deficient in cytochrome *c* and associated biogenesis factors that rely on the MIA40/ERV1 disulfide relay for import into the IMS. Taken together, our results reveal the contribution of impaired bioenergetics and mtISR signaling for mitochondrial homeostasis and cardiac health, which are compromised by CHCHD10 insolubility.

## Results

### CHCHD10 insolubility triggers ISR activation and cardiac remodeling in *Chchd10* mice

The dominant pathogenic S59L variant in *CHCHD10* responsible for multisystemic mitochondrial dysfunction in humans^14^ promotes CHCHD10 protein insolubility and compromise cardiac function in heterozygous *Chchd10^S55L/+^* knock-in mice (henceforth *Chchd10* mice), leading to heart failure and death by 1 year (Figure 1a)^25,26^. We confirmed these findings in rederived *Chchd10* mutant maintained on a C57Bl6/N background, whose cardiac function we characterized by longitudinal echocardiography (Figure 1b, c, S1a, b). While the lifespans of mutant mice were shortened to similar degrees in males and females (Figure S1c), we observed differences in cardiac dysfunction at 14 weeks of age: only male mutant mice showed reduced %LVEF (Figure 1b, c). In line, the cardiac dysfunction biomarker NPPA was upregulated 5.6-fold in males but only 4-fold in females *Chchd10* mice (Figure 1d), suggesting that onset of cardiac dysfunction is subject to biological sex. On the other hand, CHCHD10 protein accumulation in cardiac lysates were found to be increased to similar levels between male and female *Chchd10* mutant mice (Figure 1e) and alkaline sodium carbonate (Na_2_CO_3_) extraction studies performed on isolated cardiac mitochondria showed reduced solubility of CHCHD10 in mutant mice of both sexes, both at pH9.5 and pH11.5 (Figure 1f). The increased abundance of CHCHD10 in the insoluble pellet fraction in *Chchd10* mutant cardiac mitochondria did not reflect a general disruption of protein insolubility, as other mitochondrial markers such as MT-CO2, ANT1, and HSP60 behaved similarly between genotypes (Figure 1f). CHCHD10 insolubility preceded the onset of cardiomyopathy in both male (Figure 1d) and female mice (Figure 1f), and was associated with an induction of the ISR, which we could measure by cardiac bulk RNAseq (Figure 1g, Supplemental Dataset 1) and qRT-PCR analyses of marker genes *Atf4*, *Atf5*, *Trib3*, *Mthfd2*, *Phdgh*, *Asns, Aldh18a1,* and *Fgf21* (Figure S1e). Transcriptomic analyses of differentially expressed genes (DEGs) revealed that induction of core ISR gene expression signatures^36^ were similar between male and female *Chchd10* mutant mice when compared to sex-matched wild type (WT) controls (Figure 1g. S1e). Taken together, our data support the model^21^ that the earliest defects leading to CHCHD10 insolubility and ISR induction trigger the onset of cardiac dysfunction.

**Figure 1.**
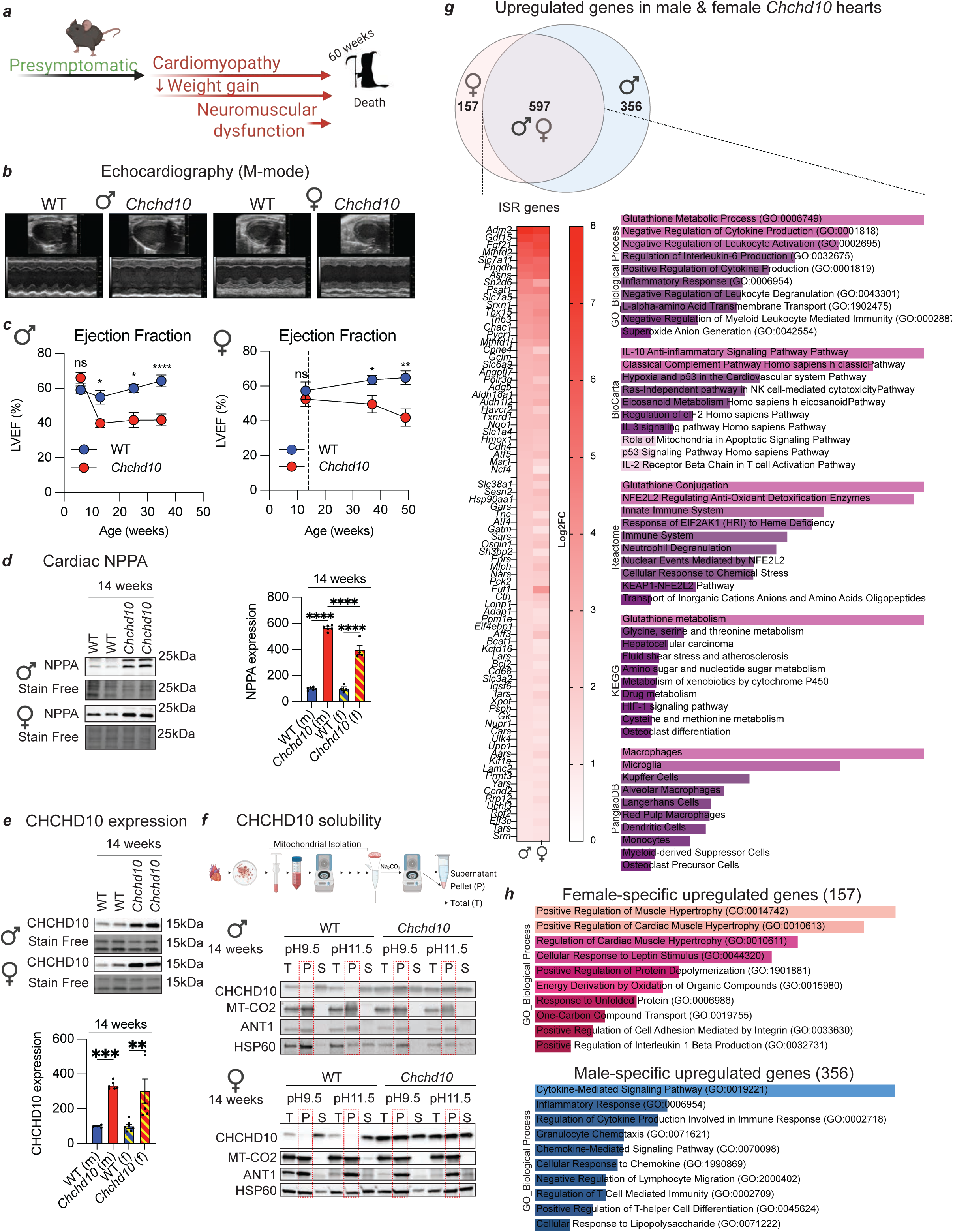
– CHCHD10 insolubility triggers ISR activation and cardiac remodeling. a) *Chchd10* heterozygous missense Ser55L (S55L) mutation causes tissue-specific defects in mice and reduced lifespan in *Chchd10^S55L/+^* mutant mice. The presymptomatic-symptomatic transition to cardiomyopathy precedes neuromuscular dysfunction. Created with Biorender.com b) Representative M-Mode echocardiographic images of the left ventricle of wild type (WT) and *Chchd10* male and female mice at 14 weeks of age. c) Left ventricular ejection fraction (% LVEF) of WT (blue, n=3-9) and *Chchd10* (red, n=3-8) male mice (left) and WT (blue, n=4-9) and *Chchd10* (red, n=6-7) female mice (right). Data represent mean *±* SEM. Dotted line represents 14 weeks. One-way ANOVA, * P<0.05, **P<0.01, ****P<0.0001, ns=not significant. d) Representative immunoblots of NPPA protein levels in cardiac lysates of male and female mice at 14 weeks of age. Densitometric quantification of WT (blue, male n=6, female n=4) and *Chchd10* (red, male n=6, female n=4) mice is relative to stain-free. Data are means ± SEM, 2-tailed unpaired Student’s t test. ****P<0.0001. e) Representative immunoblots of CHCHD10 protein accumulation in cardiac lysates of male and female mice at 14 weeks of age. Densitometric quantification of WT (blue, male n=6, female n=7) and *Chchd10* (red, male n=6, female n=6) mice is relative to stain-free. Data are means ± SEM, 2-tailed unpaired Student’s t test. **P<0.01, ***P<0.001. f) Alkaline carbonate (Na_2_CO_3_) extraction of cardiac mitochondria performed on WT and *Chchd10* mutant male (top) and female (bottom) mice at 14 weeks of age. Total (T), insoluble pellet (P), and soluble supernatant (S) fractions were analyzed by immunoblotting with indicated antibodies. Dotted outline of pellet fraction. g) Venn diagram and heatmap of upregulated differentially expressed genes (DEGs) in *Chchd10* mice. Bulk RNA-seq was performed on cardiac biopsies from male (n=3) and female (n=3) *Chchd10* mice compared to sex-matched littermate controls at 14 weeks of age. The Venn diagram shows 597 overlapping (purple), 157 female-specific (pink), and 356 male-specific (blue) upregulated DEGs. Integrated stress response (ISR) genes^125^ were significantly upregulated in both sexes (heatmap), and pathway enrichment analysis using Gene Ontology (GO), BioCarta, Reactome, KEGG, and PanglaoDB databases confirmed ISR pathway enrichment among these DEGs. h) Gene Ontology (GO) pathway analyses of female-specific upregulated gene (157) and male-specific upregulated genes (356) identified by bulk RNAseq in Figure 1g.

Although transcriptomic analyses revealed the magnitude and onset of ISR induction were similar between male and female mutant hearts (Figure 1g, Supplemental Dataset 2), we observed sex-specific differential gene expression, raising the possibility that cell signaling downstream of mitochondrial dysfunction may modulate the trajectory of cardiomyopathy. The 157 female-specific upregulated DEGs in *Chchd10* mutant hearts were associated with cardioprotective pathways involving BMP10^37^ and NR4A3^38^, whose upregulation have been independently demonstrated to protect against cardiomyopathy. In male mutant mice, pathway enrichment based on the 356 upregulated male-specific DEGs revealed an induction of immune, inflammatory, and chemokine signaling characteristic of host-pathogen interaction pathways (including viruses, bacteria and parasites) (Figure 1h, Supplemental Dataset 2).

These enriched pathways also emerged when comparing all 955 upregulated DEGs in symptomatic *Chchd10* male mice to WT littermate male mice, but the same was not observed in females (Figure S1f, g, Supplemental Dataset 3, 4). Downregulated pathways including those associated with mtDNA transcription were similar enriched in both male and female *Chchd10* mice, consistent with a relative reduction of mtDNA we observed at the onset of cardiomyopathy^26^ (Figure S1h). As *Chchd10* mutant mice were housed in germ-free conditions, we wondered whether overactive innate immune signaling triggered in response to mitochondrial dysfunction may contribute to cardiac inflammation and dysfunction as in other models of MD^39,40^. To test this hypothesis, we blunted innate immune signaling in *Chchd10* mutant mice via the whole-body ablation of Stimulator of interferon genes (STING) by crossing *Chchd10* mice with the Goldenticket mouse (*Sting^Gt/Gt^*, Figure S1i), which carries a loss-of-function I199N mutation in *Sting,* effectively knocking out the gene^41^. *Sting* encodes a transmembrane ER protein that acts as an adaptor protein involved in interferon signaling that can be triggered by microbial infection and mitochondrial dysfunction to activate innate immunity^42^. Echo analyses of *Chchd10^S55L/+^Sting^Gt/Gt^* (henceforth C*hchd10/Sting*) mutant mice revealed rescued cardiac function at 38 weeks but impaired %LVEF at 48 weeks of age (Figure S1j), pointing to a temporary, cardioprotective effect of inhibiting overactive innate immune signaling downstream of mitochondrial dysfunction. *Sting* deletion did not negatively impact cardiac function in *Sting^Gt/Gt^* (henceforth *Sting*) mice (Figure S1j) and did not influence the reduced body mass or lifespan shortening in *Chchd10/Sting* mice (Figure S1k, l). Taken together, our data indicate that inflammatory signaling triggered by mitochondrial dysfunction can modulate the progression of cardiac dysfunction downstream caused by mutant CHCHD10 insolubility.

### Impaired mitochondrial respiration is an early defect in mutant *Chchd10* hearts

Mutant CHCHD10 triggers ISR induction and subsequently the metabolic rewiring of iron-dependent and iron-sulfur cluster (ISC) pathways that are required for OXPHOS function^21^. Since OXPHOS dysfunction was observed in late-stage *Chchd10* mice, this led to the notion that impaired mitochondrial respiration is a consequence rather than a cause of cardiomyopathy^21^. When we measured oxygen consumption rates by high-resolution respirometry in cardiac mitochondria isolated from *Chchd10* mutant mice beginning at 14 weeks of age, we observed oxygen consumption defects on both carbohydrate and fatty acid-derived substrates (Figure 2a,b), which were equivalently reduced in both (symptomatic) male and (pre-symptomatic) female mutant mice. These defects were not accompanied by loss of mitochondrial membrane potential, arguing against mitochondrial uncoupling (Figure 2c, d). We observed normal mitochondria respiration from livers of *Chchd10* mice, in which CHCHD10 solubility is unaffected (Figure 2e, f), further strengthening the association between CHCHD10 insolubility and bioenergetic dysfunction. Mitochondrial respiration and membrane potential were unaffected in *Chchd10* cardiac mitochondria isolated from presymptomatic male mice at 7 weeks of age (Figure S2a-c). These data argue that bioenergetic impairment of mitochondria is an early dysfunction associated with CHCHD10 insolubility, which can precede or manifest coincidently with cardiac dysfunction.

**Figure 2.**
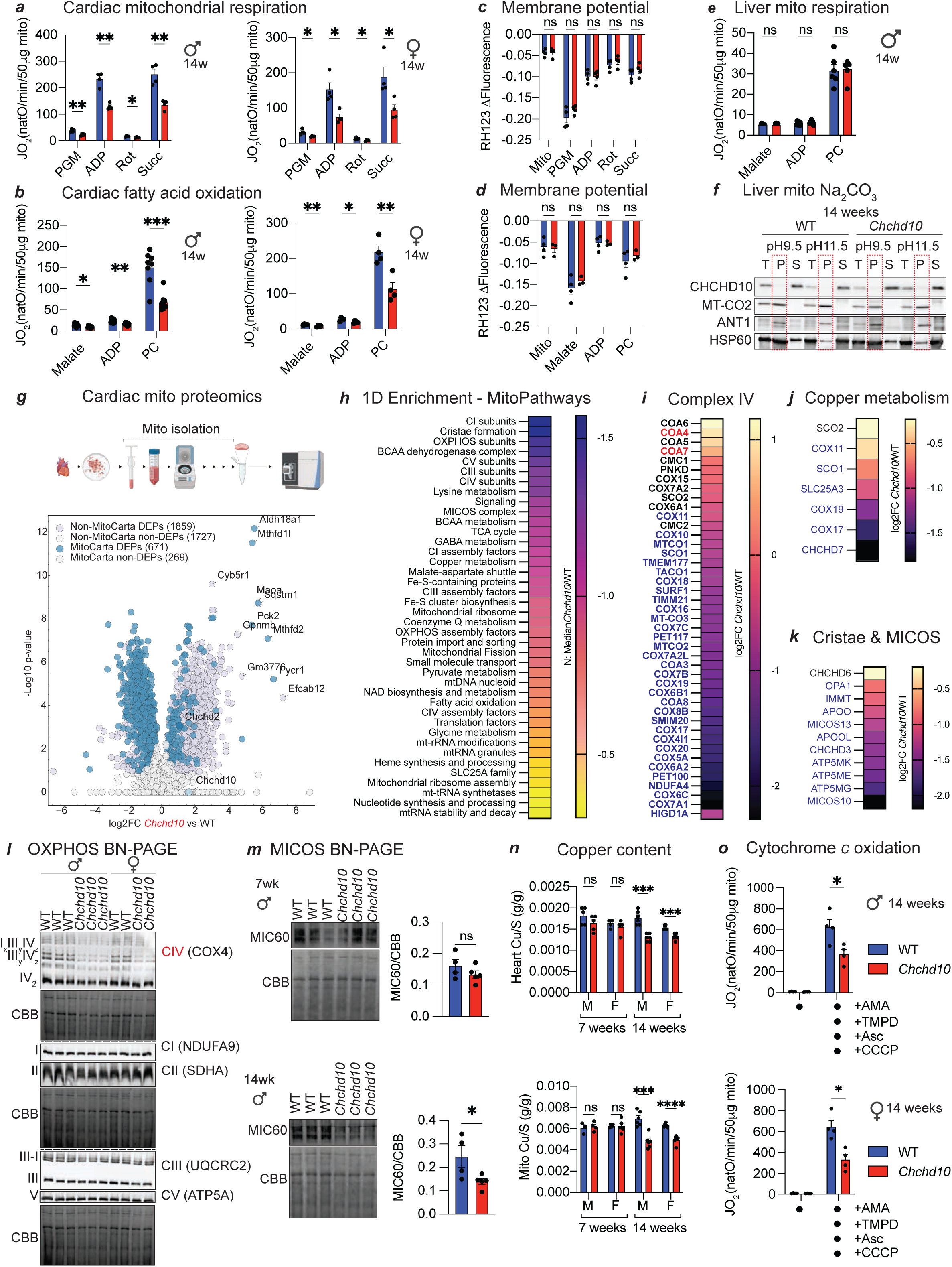
– Impaired mitochondrial respiration in mutant *Chchd10* hearts. a) Oxygen consumption rates (JO_2_) of cardiac mitochondria isolated from WT (n=4) and *Chchd10* (n=4) male (left) and female (right) mice at 14 weeks. JO_2_ measured sequentially in the presence of pyruvate, glutamate, malate (PGM), adenosine diphosphate (ADP), rotenone (Rot), and succinate (Succ). Data represent mean ± SEM; multiple unpaired t-test, *P<0.05, **P<0.01 b) Oxygen consumption rates (JO_2_) of cardiac mitochondria isolated from WT (n=4-8) and *Chchd10* (n=4-8) male (left) and female (right) mice at 14 weeks. JO_2_ measured sequentially in the presence of malate, adenosine diphosphate (ADP), and palmitoyl carnitine (PC). Data represent mean ± SEM; multiple unpaired t-test, *P<0.05, **P<0.01, ***P<0.001. c) Mitochondrial membrane potential (ΔΨ) measured by quenching of Rhodamine 123 (RH123) fluorescence in cardiac mitochondria of WT (n=4) and *Chchd10* (n=4) female mice from Figure 2a (right). Data represent mean ± SEM; multiple unpaired t-test, ns=not significant. d) Mitochondrial membrane potential (ΔΨ) measured by quenching of Rhodamine 123 (RH123) fluorescence in cardiac mitochondria of WT (n=4) and *Chchd10* (n=3) male mice from Figure 2b (right). Data represent mean ± SEM; multiple unpaired t-test, ns=not significant. e) Oxygen consumption rates (JO_2_) of liver mitochondria isolated from WT (n=6) and *Chchd10* (n=6) male mice at 14 weeks. JO_2_ measured sequentially in the presence of malate, adenosine diphosphate (ADP), and palmitoyl carnitine (PC). Data represent mean ± SEM; multiple unpaired t-test, ns=not significant f) Alkaline carbonate (Na_2_CO_3_) extraction of liver mitochondria performed on WT and *Chchd10* mutant male mice at 14 weeks of age. Total (T), insoluble pellet (P), and soluble supernatant (S) fractions generated at pH9.5 and 11.5 were analyzed by immunoblotting with indicated antibodies. Dotted outline of pellet fraction. g) Volcano plot of differentially expressed proteins (DEPs) identified by proteomics of isolated cardiac mitochondria (Supplemental Dataset 5) from WT (n=4) and *Chchd10* (red, n=4) male mice at 14 weeks of age. 671 DEPs belonging to MitoCarta 3.0 were identified. Horizontal dotted line represents –log_10_(padj)>0.05. h) 1D enrichment analysis of MitoPathways (MitoCarta 3.0) revealing the dysregulated pathways in Figure 2g (Supplemental Dataset 5). i) Heatmap of Complex IV proteins and assembly factors that were quantified by cardiac proteomics in Figure 2g and significantly upregulated (red), downregulated (blue) or unchanged (black) in *Chchd10* cardiac mitochondria relative to wild type. j) Heatmap of Copper metabolism proteins that were quantified by proteomics in Figure 2g and significantly downregulated (blue) or unchanged (black) in *Chchd10* cardiac mitochondria relative to wild type (Supplemental Dataset 5). k) Heatmap of Cristae and MICOS complex proteins that were quantified by proteomics in Figure 2g and significantly downregulated (blue) or unchanged (black) in *Chchd10* cardiac mitochondria relative to wild type (Supplemental Dataset 5). l) BN-PAGE immunoblot analysis of cardiac OXPHOS complexes isolated from WT and *Chchd10* male and female mice at 14 weeks using the indicated antibodies for Complex IV (COX4), Complex I (NDUFA9), Complex II (SDHA), Complex III (UQCRC2) and Complex V (ATP5A). Coomassie brilliant blue (CBB) was used as loading control. Quantification in Figure S2f. m) Representative BN-PAGE immunoblot analysis of cardiac MICOS complexes isolated from WT and *Chchd10* male mice at 7 and 14 weeks using the indicated antibody against IMMT/MIC60. Densitometric quantification is relative to Coomassie brilliant blue (CBB). Data are means ± SEM, 2-tailed unpaired Student’s t test. * P<0.05, ns=not significant. n) Copper content in total heart (top, n=5-6) and cardiac mitochondria (bottom, n=3-5) samples measured by inductively coupled plasma–optical emission spectrometry (ICP-OES) from wild type (WT) and *Chchd10* male (M) and female (F) mice at 7 and 14 weeks of age. Data represent mean values normalized to sulfur (S) *±* SEM, Multiple 2-tailed unpaired Student’s t test, ***P<0.001, ****P<0.0001, ns=not significant. o) Oxygen consumption rates (JO_2_) of cardiac mitochondria from WT (n=4) and *Chchd10* (n=4) male (top) and female (bottom) mice at 14 weeks incubated with antimycin A (AMA) to prevent electron transfer from Complex III followed by addition of carbonyl cyanide m-chlorophenyl hydrazine (CCCP), and N,N,N′,N′-Tetramethyl-p-phenylenediamine (TMPD) and ascorbate (Asc) to measure cytochrome *c* oxidation. Data represent mean ± SEM; unpaired Student’s t-test, *P<0.05

To understand why mitochondrial respiration was reduced in response to mutant CHCHD10, we compared the proteomes of cardiac mitochondria isolated from WT and *Chchd10* mice, which revealed a dramatic remodeling: 71% of the 940 quantified mitochondrial proteins belonging to MitoCarta 3.0 were differentially expressed, with most of them being downregulated (average log2 FC = –0.7) (Figure 2g). Stratifying the MitoCarta 3.0 DEPs based on known submitochondrial localization (OMM, IMS, IMM, or matrix) did not reveal submitochondrial compartment biases, highlighting a generally uniform dysregulation of mitochondrial proteostasis (Figure S2d). One-dimensional enrichment analyses revealed several MitoPathways, notably those implicated in the maintenance and assembly of OXPHOS and MICOS complexes and copper metabolism (Figure 2h-k, S2e). Plotting quantified MitoCarta 3.0 proteins revealed significant reductions in the steady-state levels of components of Complex IV (Figure 2i) as well as other OXPHOS complexes (Figure S2e). BN-PAGE analyses revealed normal levels of Complexes I, II, III, and V and a specific reduction in Complex IV assemblies (Figure 2l, S2f), which accompanied the reduced levels of subunits and assembly factors such as COA8, COX16, COX18, COX20, COX4L1, COX5A, COX6B1, COX6C, COX7A1, COX7A2L, COX7B, COX7C, COX8B, COX10, COX15, COX18, COX20, and the mtDNA-encoded proteins MT-CO1, MT-CO2 and MT-CO3. We observed a reduction in HIGD1A (Figure 2i), which coordinates the assembly of COX-containing complexes^43^, and a shift in the balance between COX6A1 and COX6A2 isoforms, which were reported to impact the stability of COX-containing supercomplexes^44^. High resolution fluor-respirometry performed in isolated cardiac mitochondria in the presence of Antimycin A, TMPD, Ascorbate, and CCCP, which is typically used to measure Complex IV activity^45^, revealed a ∼ 40% decrease in oxygen consumption rates (JO_2_) in both (symptomatic) male and (presymptomatic) female mice at 14 weeks (Figure 2o), pointing to a defect in cytochrome *c* oxidation. We observed reductions in copper handling proteins COX11, COX19, SCO1, CHCHD7, and the IMM copper transporter SLC25A3 (Figure 2j), all of which are required for the metalation of COX^46^. In addition, COX17, which supplies the Cu necessary via SOC1 to assemble both the COX1 and COX2 modules required to assemble the COX holoenzyme, was also reduced. In yeast, COX17 is involved in the assembly of COX as well as the MICOS complex^47^, an integral membrane complex that bridges outer and inner membranes and is required to maintain cristae structure^48^. Proteomic profiling of isolated cardiac mitochondria also showed a reduction in MICOS subunits MIC60/IMMT, MIC13, CHCHD3, APOO, and APOOL and the MICOS-interactor OPA1 (Figure 2k), which also plays a central role in the maintenance of cristae structure^49^. In line, BN-PAGE analysis of the MICOS complex revealed a reduction in MIC60 immunoreactivity at 14 weeks (but not 7 weeks) of age (Figure 2m), which paralleled the disruption previously reported in *CHCHD10^S59L/+^* patient-derived fibroblasts^20^.

Copper is an essential redox cofactor for several mitochondrial enzymes, including Cytochrome *c* oxidase (Complex IV), whose metalation in the CuA and CuB sites is essential for assembly and enzymatic activity^46^. As genetic defects in disrupting the insertion of copper into Complex IV cause severe, multisystemic defects including heart failure in mice and humans^50–55^, we measured copper levels in *Chchd10* mutant hearts by inductively coupled plasma optical emission spectroscopy (ICP-OES), which enables the direct, precise, sensitive, and accurate measurement of metals in biology^56^. ICP-OES analyses uncovered a reduction in cardiac and mitochondrial (Figure 2n) copper content in 14-week-old mutant male and female mice, which paralleled the observed reduction in cytochrome *c* oxidation rates (Figure 2o). In contrast to previous studies^21^, no reductions in iron nor heme were observed at either 7 or 14 weeks of age (Figure S2g, h). ICP-OES revealed other metals such as zinc (Zn), magnesium (Mg), or manganese (Mn) were unaltered in *Chchd10* hearts and cardiac mitochondria (Figure S2i). Altogether, our data reveal that a defect in CHCHD10 protein solubility is associated with impaired cytochrome *c* oxidation, which may contribute to the pathological cardiac remodeling in *Chchd10* mutant mice.

### Inhibition of OMA1 catalytic activity suppresses mtISR but not CHCHD10 insolubility

Previous studies of *Chchd10* mice have intimated that the mtISR induction is responsible for OXPHOS dysfunction via the maladaptive rewiring of mitochondrial metabolism^21,25,28^. CHCHD10^S59L^ triggers the activation of the stress-induced metalloprotease OMA1, which proteolytically processes DELE1 in the IMS so that the cleaved form can be exported to the cytosol where it signals through the heme-regulated inhibitor (HRI) kinase to activate the ISR^8,9,19,57,58^. Monitoring the proteolytic cleavage of another classical OMA1 substrate L-OPA1 revealed OMA1 activation in symptomatic *Chchd10* mutant mice (14 weeks of age) that was limited in presymptomatic hearts (7 weeks of age), which showed elevated levels of MTHFD2, SQSTM1/P62, and LC3-II (Figure S3a), indicating that stress-induced OPA1 processing occurs after ISR induction. Since OMA1 ablation can suppress cardiac dysfunction and mitochondrial fragmentation in the hearts of cardiomyocyte-specific *Yme1l1* knockout mice (*Yme1l1^Heart^*), which also show increased OMA1 activity^59^ and ISR signaling according to bulk RNAseq studies performed at 35 weeks of age (Figure S3b), we wondered whether inactivation of OMA1 could confer cardioprotection to *Chchd10* mutant mice. Whole-body deletion of *Oma1* is synthetically lethal in *Chchd10* mutant mice carrying the G58R variant^19^, prompting us to generate a catalytic site mutant E324Q in OMA1 via Crispr/Cas9 genome editing of C57Bl/6N mice (Figure 3a, b), which is known to inhibit the proteolytic activity of OMA1 without disrupting putative, non-catalytic scaffolding functions in the IMM^60,61^. *Oma1^E324Q/E324Q^*(henceforth *Oma1*) mutant mice were outwardly normal and cardiac transcriptomic profiling performed at 14 weeks of age revealed virtually no gene dysregulation: 6 and 3 out of 17,1911 genes were differentially expressed in *Oma1* male and female mice, respectively (Figure S3c, Supplemental Dataset 1). Similarly, proteomic comparisons of isolated cardiac mitochondria from WT and *Oma1* male mice revealed no differentially expressed proteins (DEPs) (Figure S3d, Supplemental Dataset 5), consistent with previous proteomics analyses of cardiomyocyte-specific *Oma1* knockout mice^31^. We confirmed the catalytic inactivation of OMA1 in mouse fibroblasts derived from *Oma1* embryos by examining constitutive and CCCP-induced OPA1 processing (Figure S3e). Next, we intercrossed *Oma1* and *Chchd10* mutant mice and successfully generated *Oma1^E324Q/E324Q^Chchd10^S55L/+^* (*Chchd10/Oma1*) double mutant mice at Mendelian ratios (Figure 3b). Double mutant male and female mice, which were outwardly normal, showed a marked reduction in cardiac ISR induction by RT-qPCR, and bulk RNAseq analyses (Figure 3c, S3f, g, Supplemental Dataset 1,6), further validating the suppression of OMA1 activity and the mtISR in these mice. Similarly, proteomic profiling in male mice revealed a reduction in ISR protein levels in *Chchd10/Oma1* isolated cardiac mitochondria in comparison to those isolated from *Chchd10* mice (Figure 3d). Most of the upregulated DEPs that were suppressed in *Chchd10/Oma1* hearts (relative to *Chchd10*) were DELE1-dependent mtISR factors^22,36^ including PYCR1, AKR1B7, MTHFD2, PCK2, MTHFD1L, ALDH18A1, GHITM, GPT2, LONP1, GARS1, GATM, and SHMT2 (Figure S3h). However, suppression of mtISR signaling was not associated with rescued solubility of mutant CHCHD10: Na_2_CO_3_ extraction studies revealed CHCHD10 to be equally insoluble in *Chchd10* and *Chchd10/Oma1* relative to WT and *Oma1* cardiac mitochondria (Figure 3e), demonstrating that CHCHD10 insolubility can be uncoupled from mtISR induction.

**Figure 3.**
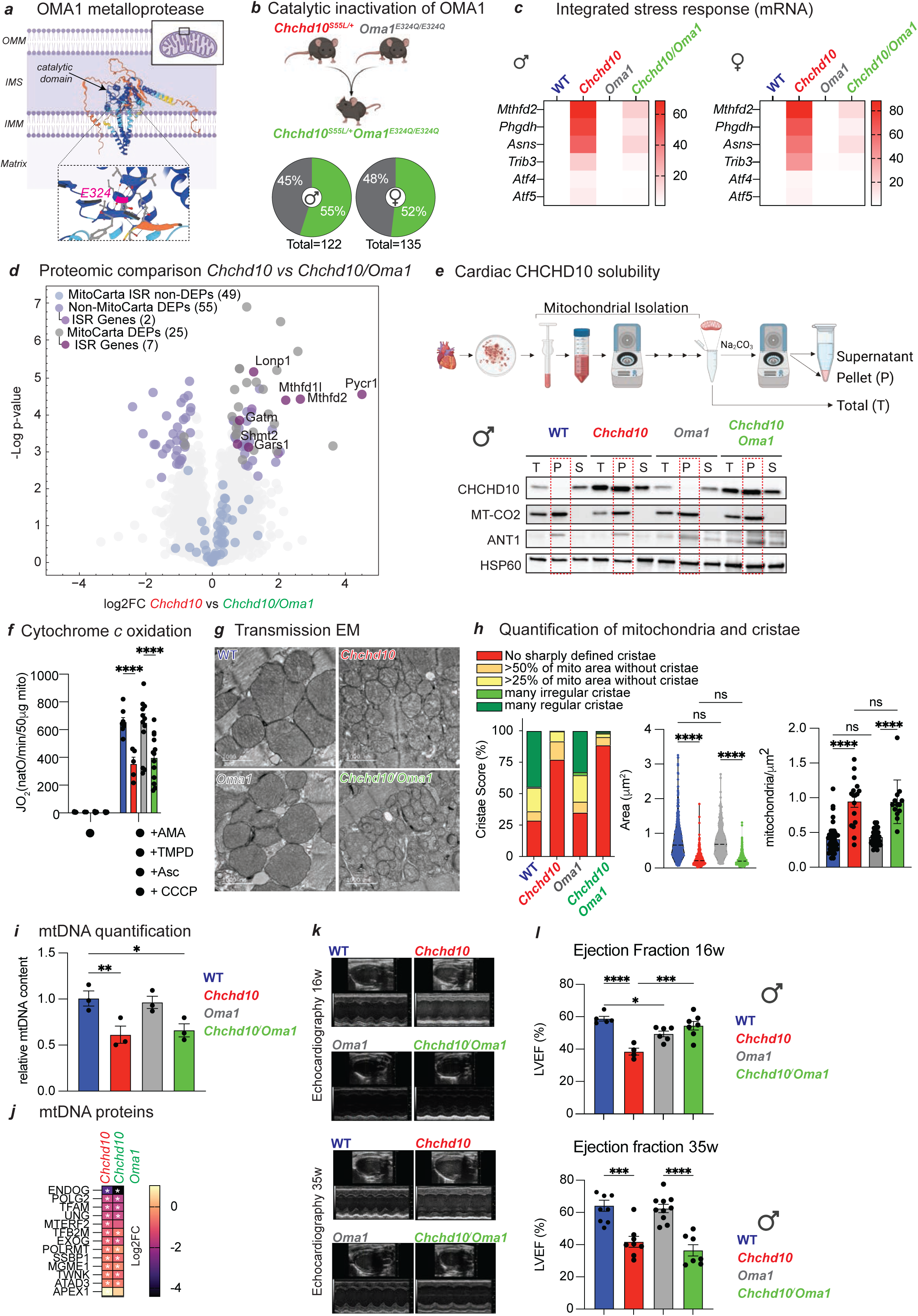
– *Oma1^E324Q/E324Q^* mice suppress mtISR and bypass mitochondrial dysfunction. a) Structural representation of OMA1 metalloprotease with Alphafold 3.0. Catalytic core including the E324 glutamic acid residue within the conserved HEXXH motif that is essential for zinc binding and catalytic activity is indicated in the inset. Substitution to glutamine (E324Q) inhibits catalytic activity. Created with Biorender.com b) Generation of the *Chchd10^S55L/+^Oma1^E324Q/E324Q^* (*Chchd10/Oma1,* green) mice by intercrossing *Chchd10^S55L/+^* (*Chchd10*, red) with *Oma1^E324Q/E324Q^* (*Oma1*, grey) mice. Generation of male (n=122) and female (n=135) offspring from *Chchd10/Oma1* intercrosses with *Oma1* mice according to Mendelian ratios: chi-squared for male 1.18<3.84 (p=0.05, df=1) and female 0.185<3.84 (p=0.05, df=1) mice. Created with Biorender.com c) Heat map of integrated stress response (ISR) genes expression measured by qRT-PCR of total cardiac biopsies from wild type (blue, n=3), *Chchd10* (red, n=3), *Oma1* (grey, n=3), and *Chchd10/Oma1* (green, n=3) male and female mice at 14 weeks (see Figure S3f, g for one-way ANOVA). d) Volcano plot of cardiac proteomics performed on cardiac mitochondria isolated from male WT (n=5) and *Chchd10* (n=5) mutant male mice at 14 weeks of age. Proteins ascribed to MitoCarta 3.0 and integrated stress response (ISR) genes are indicated. Differentially expressed proteins (DEPs) plotted according to the log2 fold change of *Chchd10* vs *Chchd10/Oma1* versus the –log10 transformed p-value of a two-sided t-test. Significance was considered for a permutation based-FDR cutoff of 0.05. Mitochondrial (MitoCarta 3.0) and ISR Genes as previously defined^126^ are highlighted by color. e) Alkaline carbonate (Na_2_CO_3_) extraction of cardiac mitochondria performed on cardiac mitochondria isolated from wild type (WT), *Chchd10*, *Oma1*, and *Chchd10/Oma1* mice at 14 weeks of age. Total (T), insoluble pellet (P), and soluble supernatant (S) fractions were analyzed by immunoblotting with indicated antibodies. Dotted outline of pellet fraction. f) Oxygen consumption rates (JO_2_) of cardiac mitochondria from male WT (blue, n=8), *Chchd10* (red, n=5), *Oma1* (grey, n=13), and *Chchd10/Oma1* (green, n=13) male mice at 14 weeks incubated with antimycin A (AMA) to prevent electron transfer form Complex III followed by addition of carbonyl cyanide m-chlorophenyl hydrazine (CCCP), and N,N,N′,N′-Tetramethyl-p-phenylenediamine (TMPD) and ascorbate (Asc) to measure cytochrome *c* oxidation. Data represent mean ± SEM; unpaired Student’s t-test, ****P<0.0001 g) Representative transmission electron micrographs (EM) of cardiac posterior walls of WT (blue, n=3) and *Chchd10* (red, n=3) *Oma1* (grey, n=3), and *Chchd10/Oma1* (green, n3) male mice at 14 weeks. Scale bar: 1000 nm. h) Quantification of Figure 3g performed with Image J to assess the area of manually-segmented mitochondria area (mm^2^) and the number of mitochondria per image area (mitochondria/mm^2^) from WT (blue, n=3), *Chchd10* (red, n=3), *Oma1* (grey, n=3), and *Chchd10/Oma1* (green, n=3) mice at 14 weeks. 315-440 mitochondria were quantified per condition. Cristae score was assigned blindly according to previously defined classification^129^. Data are means ± SEM, 2-tailed unpaired Student’s t test. ****P<0.0001. Thick dotted lines represent median values. i) Quantification of mitochondrial DNA (mtDNA) in cardiac biopsies from male WT (blue, n=3), *Chchd10* (red, n=3), *Oma1* (grey, n=3), and *Chchd10/Oma1* (green, n=3) mice at 14 weeks. Primers directed at mtDNA encoding 16s rRNA and β-actin for nDNA were used. Data are means ± SEM, One-way ANOVA, * P<0.05, ** P<0.01. j) Heat map of significantly downregulated (*) mtDNA maintenance proteins in cardiac mitochondria profiled from *Chchd10* (red, n=5) versus WT (n=5) and *Chchd10/Oma1* (green, n=5) vs WT hearts by mass spectrometry (Supplemental Dataset 5). k) Representative M-Mode echocardiographic images of the left ventricle of wild type (WT), *Chchd10*, *Oma1*, and *Chchd10/Oma1* male mice at 16 weeks (top) and 35 weeks (bottom) of age. l) Left ventricular ejection fraction (% LVEF) of WT (blue, n=5-8), *Chchd10* (red, n=4-8), *Oma1* (grey, n=6-10), and *Chchd10/Oma1* (green, n=7) male mice at 16 weeks (top) and 35 weeks (bottom) of age. Data represent mean *±* SEM. One-way ANOVA, * P<0.05, ***P<0.001, ****P<0.0001.

### Inhibition of OMA1 delays cardiomyopathy in *Chchd10* mutant mice

Having generated viable *Chchd10* mutant mice with inactive OMA1 and blunted mtISR, we decided to explore whether OMA1 inactivation modulates mitochondrial dysfunction in *Chchd10/Oma1* hearts. High-resolution respirometry revealed that cytochrome *c* oxidation rates were equivalently diminished in male *Chchd10/Oma1* and *Chchd10* cardiac mitochondria relative to either WT and *Oma1* littermates (Figure 3f). In line, principal component analyses (PCA) of mitochondrial proteomics showed that *Chchd10* and double mutant mice overlap substantially, indicating similarity in the mitochondrial proteomes of these two groups, which were clearly demarcated from both WT and *Oma1* cardiac proteomes (Figure S3i). These data suggest that bioenergetic defects observed at the onset of cardiomyopathy in *Chchd10* mice are not caused by mtISR activation, which is consistent with previous in vitro observations in HEK293T cells expressing insoluble, mutant CHCHD10^G58R^ in which *OMA1* silencing did not rescue mitochondrial respiration^19^. TEM analyses of mitochondrial size and cristae content showed that defects in mutant *Chchd10* hearts (Figure 3g, h) were not rescued by OMA1 inactivation in *Chchd10/Oma1* hearts, despite an inhibition of stress-induced OPA1 processing. We observed a 36% reduction of mtDNA at 14 weeks in both male and female *Chchd10* mutant hearts by qPCR (Figure 3i) that mirrored the reduction in mtDNA factors such as TFAM, the mtDNA nucleoid regulator whose abundance tracks and controls mtDNA content^62–64^, as well as POLG2, POLRMT, TWINKLE, and mtSSB (Figure 3j). These mtDNA defects were not rescued in *Chchd10/Oma1* mice at 14 weeks of age (Figure 3i, j), arguing against a role of the mtISR in regulating mtDNA content at the onset of cardiac dysfunction^21^. Despite the persistence of structural and functional defects in mitochondria in the hearts of *Chchd10/Oma1* mice, echocardiography revealed an improvement of cardiac function in double mutant mice (Figure 3k, l): reduced %LVEF was restored to levels indistinguishable from wild type mice in male mutant mice at 16 weeks of age (Figure 3l) pointing to a cardioprotective effect of OMA1 inactivation. OMA1 inactivation did not rescue cardiac fibrosis induced by mutant CHCHD10 (Figure S3j) and by 35 weeks of age, cardiac function in double mutant mice declined to levels observed in *Chchd10* mutant mice, indicating that OMA1-dependent cardioprotection is temporary (Figure 3k,l). Of note, both male and female double mutant male and female mice had lifespans and weight gain curves that were similar to *Chchd10* mutant counterparts, suggesting that the molecular mechanisms through which OMA1 inhibition influences organ function may be tissue-specific^19,22^ (Figure S3k, l). Taken together, our data suggests that OMA1 inactivation delays the onset of cardiomyopathy independently of the modulation of cytochrome *c* oxidation, mtDNA content, and cristae structure defects in mutant *Chchd10* mice.

To gain insights into how OMA1 inactivation modulates downstream cardiac signaling we analyzed cardiac transcriptomes by bulk RNAseq. First, we decided to compare female *Chchd10/Oma1* to female *Chchd10* mice since both have normal cardiac function at 14 weeks, enabling us to identify OMA1-specific pathways that can be modulated in response to mutant CHCHD10. In line with a reduction of ISR signaling, we observed 31 out 67 DEGs in *Chchd10/Oma1* double mutant hearts were involved in ATF-dependent signaling (Figure S3m, Supplemental Dataset 1), consistent with qRT-PCR studies revealing a reduction in ISR markers (Figure 3c). Consequently, Enrichr pathway analyses of the 31 upregulated DEGs in *Chchd10* hearts relative to *Chchd10/Oma1* hearts revealed an implication of amino acid metabolism (glycine, serine, threonine, cysteine and methionine), ferroptosis, and folate and one-carbon metabolism (Figure S3m, Supplemental Dataset 6) and stress signaling pathways associated with PERK, GCN2, ATF and HRI signaling. On the other hand, 55 genes were upregulated in *Chchd10/Oma1* hearts relative to *Chchd10* hearts, which included factors involved in anti-viral and interferon signaling pathways such as *Cxcl9, Cxcl10*, *Cxcl12*, and *Cxcl21a* as well as *Irf7*, *Isg15*, *Oasl1*, *Irgm1*, and *Mx2*. Closer inspection revealed a peculiar albeit limited set of upregulated genes belonging to these clusters, which included IFIT family members (*Ifit1*, *Ifit2*, *Ifit3, and Ifit3b*), which have previously been identified as negative regulators of pathogen-induced NF-kappaB and TNF signaling^65–67^. In line, analysis of sirius red cardiac histology at 22 weeks of age revealed a reduction in cardiac fibrosis in female *Chchd10/Oma1* mice relative to female *Chchd10* mice, consistent with a cardioprotective role of OMA1 inactivation. In males, cardiac fibrosis was equally elevated in both *Chchd10/Oma1* and *Chchd10* mice at 22 weeks of age. The degree of cardiac fibrosis in male and female *Chchd10* mice showed no sexual dimorphism, suggesting that female hearts may cope better that male hearts in response to mitochondrial dysfunction (Figure S3j-l). Indeed, the comparison of DEGs in WT male and female littermates at 14 weeks of age revealed a male-specific upregulation of inflammatory factors *C7* and *Ccl11*, as well as *Nppb* (Supplemental Dataset 1), which encodes the Natriuretic Peptide B associated with decreased cardiac output and increased cardiac damage^68^ that is consistent with an established, underlying sensitivity of male mice for cardiomyopathy^69^. Altogether, our data indicate that inactivation of OMA1 confers cardioprotection via transcriptional remodeling associated with suppression of the mtISR.

### CHCHD10 insolubility disrupts IMS and IMM proteostasis

Having identified downstream signaling pathways triggered by mitochondrial dysfunction in the hearts of *Chchd10* mutant mice, we focused our attention back to the molecular defects underpinning cytochrome *c* oxidation. At first, the most parsimonious explanation was an enzymatic defect in Complex IV, which was associated with reduced copper and Complex IV supercomplex levels (Figure 2l). Yet, the approximately 50% reduction in cytochrome *c* oxidation rates measured by high resolution respirometry was associated with a more modest 31% and 21% decrease in mitochondrial copper content in males and females, respectively (Figure 2n). Similarly, recent studies have called into question the functional relevance of impaired respiratory chain supercomplex (RCS) assembly for cardiac homeostasis under basal conditions^70^. Cardiac mitochondrial proteomics revealed that most DEPs associated to Complex IV were reduced (Figure 2g, i), yet pairwise comparisons of *Chchd10* and WT hearts revealed an unexpected increase in the levels of COA4 and COA7, both of which are twin Cx(9)C motif IMS proteins that are functionally linked to Complex IV via the handling of mitochondrial copper^71,72^ (Figure 2i). As the IMS import, folding, and activity of these proteins is reliant on MIA40/CHCHD4 pathway and given that MIA40/CHCHD4 levels were also found to be increased in *Chchd10* mutant mice (Figure 2g, Supplemental Dataset 5), we wondered if CHCHD10 insolubility might influence the insolubility other mitochondrial proteins relevant for cardiac health. Therefore, we performed differential solubility proteomics on *Chchd10* cardiac mitochondria. Solubilizing cardiac mitochondrial lysates in detergent followed by differential centrifugation enabled us to separate detergent-soluble supernatant fractions from detergent-insoluble pellet fractions. These fractions along with the initial non-fractionated total input were subsequently analyzed by mass spectrometry (Figure 4a). PCA distinguished WT and *Chchd10* total and supernatant fractions based on genotype, but not the pellet fraction (Figure 4b). Therefore, to identify candidate insoluble candidate proteins, we plotted mitochondrial proteins in the insoluble pellet fraction relative to their abundance measured in the total input of *Chchd10* mutant mice, which allowed us to correct for relative decreases of individual mitochondrial proteins and avoid that highly upregulated proteins be improperly ascribed to the insoluble pellet fraction simply based on their altered expression in the total input (Figure 4c). In so doing, we observed a wide range of proteins from various mitochondrial subcompartments whose relative solubility were reduced using this metric (Figure 4c), including MTFP1 and cytochrome *c*. MTFP1 is an IMM protein that regulates inner membrane integrity and bioenergetic efficiency of cardiomyocytes^73^. While it essential for adult cardiac function its ablation in cardiomyocytes does not phenocopy the bioenergetic mitochondrial defects nor the cardiac defects observed in *Chchd10* hearts^73^. Cytochrome *c* is a soluble, dual-function protein that acts as an electron donor to Complex IV at the IMM but also triggers caspase activation and apoptosis when it is released from mitochondria to the cytosol^74^. Total cytochrome *c* levels were reduced in cardiac mitochondria from *Chchd10* mutant mice by 34-fold and were increased in the insoluble pellet fraction by 34-fold. To determine whether cytochrome *c* dysfunction contributed to the reduced oxygen consumption rates we observed in isolated cardiac mitochondria supplied with Antimycin A, TMPD, Ascorbate, and CCCP (Figure 2o, 3f), we repeated high resolution respirometry measurements with mitoplasts generated from wild type and *Chchd10* cardiac mitochondria to which TMDP, and Ascorbate were added, observing once again a significant impairment of oxygen consumption rates. Subsequent addition of exogenous bovine cytochrome *c* to immediately increased oxygen consumption rates in wild type and *Chchd10* mitoplasts alike and to statistically indistinguishable levels (Figure 4d), arguing that cytochrome *c* deficiency in mitochondria from *Chchd10* mice is responsible for impaired cytochrome *c* oxidation, at least in isolated mitoplasts assayed in vitro. In this assay, increased oxygen flux occurred within a matter of seconds following injection of bovine cytochrome *c* into the respirometry chambers, (Figure S4a) arguing against an indirect increase due to mtDNA gene expression indirectly stimulated by cytochrome *c*. To test whether *Chchd10* mice suffer from an intrinsic cytochrome *c* deficiency, we analyzed cardiac lysates by immunoblot, which revealed a profound decrease in cytochrome *c* in both symptomatic and presymptomatic mice (Figure 4e) that paralleled the kinetics of CHCHD10 insolubility and accumulation and ISR signaling (Figure S3a). Transcriptomic analyses of *Chchd10* in mutant hearts did not reveal a decrease in cytochrome *c* mRNA levels (Supplemental Dataset 1) that could explain reduced steady-state protein levels, instead implying defects in the import and/or stability of cytochrome *c*.

**Figure 4.**
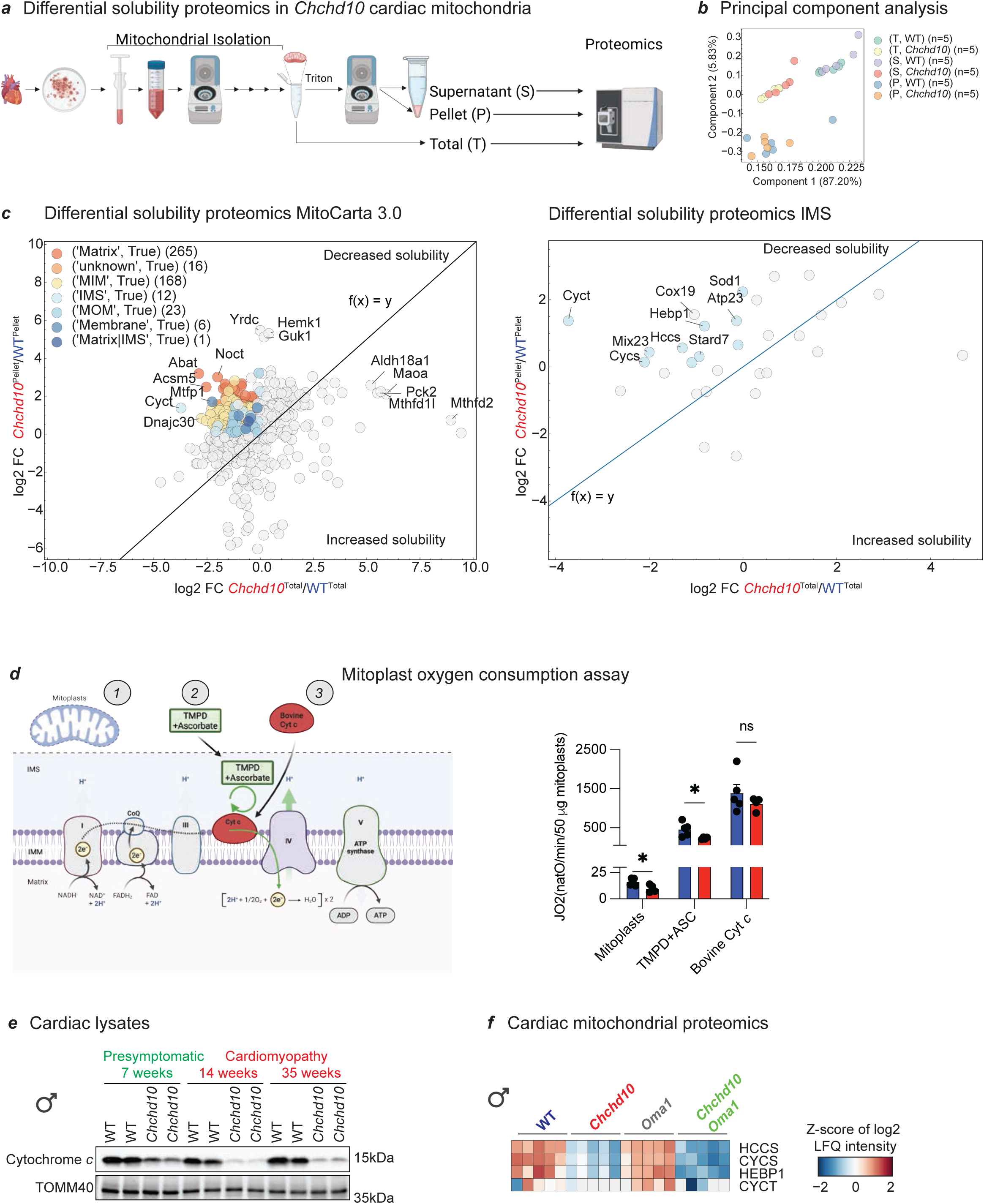
– Differential solubility proteomics identifies cytochrome *c* defect. a) Differential solubility proteomic analyses of supernatant (S) and pellet (P) fractions of detergent-solubilized cardiac mitochondria from WT and *Chchd10* male mice at 14 weeks of age. b) Principal component analysis (PCA) of Total (T) input, soluble supernatant (S), and pellet (P) fractions. Experiment was performed on 5 biological replicates. c) Mitochondrial protein insolubility assessed by measuring protein abundance (Log2FC) measured in the *Chchd10* mutant pellet (P) relative to the WT pellet (y axis) plotted against the abundance (Log2FC) measured in the *Chchd10* total input (T) relative to the WT total input (x axis) for all MitoCarta 3.0 proteins (left) or IMS proteins (right). The line represents the identity function (fx) = y (e.g. slope 1). Proteins belonging to the matrix, inner membrane (MIM), intermembrane space (IMS), outer membrane (MOM) according to MitoCarta. Proteins on the left side of the diagonal reflect reduced relative solubility (Supplemental Dataset 7). d) Oxygen consumption rates (JO_2_) of mitoplasts from WT (n = 5) and *Chchd10* (n = 4) male mice at 14 weeks. Cartoon represents workflow (left). JO_2_ measured sequentially in the presence of N,N,N′,N′-Tetramethyl-p-phenylenediamine (TMPD) plus ascorbate (ASC) and bovine cytochrome *c* (Cyt *c*) (right). Data represent mean ± SEM; unpaired Student’s t-test, * P<0.05, ns=not significant. Created with Biorender.com e) Immunoblots in cardiac lysates of presymptomatic (green) and symptomatic (red) WT and *Chchd10* male mice. Antibodies directed at cytochrome *c* and TOMM40. f) Heatmaps representing cardiac proteomic measurements of Cytochrome c (CYTC and CYCS), HEBP1, and HCCS in the total fraction of wild type (WT, n=5), *Chchd10* (n=5), *Oma1* (n=5) and *Chchd10/Oma1* (n=5) male mice at 14 weeks of age. The log2 LFQ intensities were Z-Score transformed. Rows were ordered using the complete method applying Euclidean distance. The dendrogram is not shown.

Cytochrome *c* is synthesized in the cytoplasm as apocytochrome c, lacking heme, and its import into the IMS is inexorably linked to the covalent attachment of heme at the CXXCH motif by the heme lyase <u>H</u>olo-<u>C</u>ytochrome <u>C</u>-type <u>S</u>ynthase (HCCS)^75^. The steady-state levels of HCCS and the heme-binding protein (HEBP1), which has been associated with mitochondrial dysfunction in rodents^76^, were reduced in *Chchd10* and *Chchd10/Oma1* cardiac mitochondria (Figure 4f) that showed reduced mitochondrial respiration rates in the presence of Antimycin A, TMPD, Ascorbate, and CCCP (Figure 3f). Filtering the differential solubility proteomic data on the basis of IMS proteins revealed that MIA40 clients such as COX19, SOD1, ATP23 and MIX23, and were disproportionally affected by mutant CHCHD10 (Figure 4c), consistent with a defect in IMS proteostasis and/or biogenesis (Figure S4b). Altogether, our data implicate defects in cytochrome *c* biogenesis in the impairment of mitochondrial respiration caused mutant CHCHD10, revealing an early bioenergetic defect contributing to the onset of cardiac dysfunction in *Chchd10* mutant mice.

## Discussion

A defining feature of the *CHCHD10^S59L^* human mutation (*Chchd10^S55L^* in mice) is its impact on the insolubility and accumulation of CHCHD10 within mitochondria, which is associated with a constellation of pleiotropic mitochondrial defects often observed in other mitochondrial disease (MD) animal models and patient-derived biopsies^19,21–23,25,26,28^. These defects include mtDNA depletion, cristae dysmorphology, mitochondrial fragmentation, impaired proteostasis, and activation of the OMA1-DELE1-HRI axis of the mitochondrial integrated stress response (mtISR). The convergence of multiple, often overlapping mitochondrial defects is a recurrent feature across diverse MD preclinical animal models and patient-derived samples, complicating both molecular diagnosis and therapeutic development. Dissecting which abnormalities represent primary pathogenic drivers versus downstream adaptations is therefore essential to establish causal mechanisms and to guide the design of mechanism-based interventions. Our study provides new insights into the mechanisms through which insoluble CHCHD10 promotes mitochondrial and cardiac dysfunction in a *Chchd10^S55L/+^* mutant mice by i) defining the functional relevance mtISR signaling through the catalytic inhibition of OMA1 and ii) uncovering a novel, reversable early-onset defect in cytochrome *c* oxidation. These findings challenge prevailing models in which mitochondrial bioenergetic collapse is considered a downstream consequence of chronic mtISR signaling^21,25^, instead positioning defective intermembrane space (IMS) proteostasis and cytochrome *c* deficiency as early and previously underappreciated triggers for pathogenic cardiac remodeling.

We adapted a strategy previously used to identify insoluble proteins resulting from the perturbation of mitochondrial proteostasis in yeast^77^ that we coupled to mass spectrometry to perform differential solubility proteomic approach, which enabled us to identify a subset of mitochondrial proteins with increased insolubility in *Chchd10* mutant mitochondria, implicating disruption of redox-regulated IMS protein import machinery governed by the MIA40/ERV1 axis (Figure 4). Among these, cytochrome *c* emerges as a critical node: it is insoluble in *Chchd10* female and male mutant hearts, and its levels are reduced in presymptomatic mutant mice as are HCCS, which is needed for cytochrome *c* biogenesis and stability^75,78,79^. Strikingly, supplementation with exogenous bovine cytochrome *c* rescues impaired oxygen consumption rates in cardiac mitoplasts isolated from *Chchd10^S55L/+^* mutant mice (Figure 4). This finding implicates disrupted cytochrome *c* biogenesis as a key contributor to mitochondrial respiratory collapse, which we show occurs at the onset of cardiomyopathy rather than as a late-stage event^21,25^. It will be important to determine whether this paradigm is also relevant for the late-onset neuromuscular defects described in *Chchd10* mice^20,26,80^.

The bioenergetic relevance of cytochrome *c* oxidation in Amyotrophic Lateral Sclerosis (ALS) was recently highlighted by studies revealing that the genetic disruption of Cytochrome *c* Oxidase (Complex IV) specifically in neurons is sufficient to recapitulate molecular, cellular, and neuromuscular features of sporadic ALS, including motor neuron death, neuroinflammation, and muscle wasting^81^. Convergent clinical and phenotypic manifestations of impaired cytochrome *c* oxidation due to independent cytochrome *c* or Complex IV defects have been documented in other forms of MD, such as microphthalmia with linear skin lesions (MLS)—a developmental syndrome characterized by defects of the neuromuscular, cardiovascular, and skin systems arising from mutations in either *HCCS* or *COX7B*, a structural subunit of Complex IV^82,83^. We suspect that other examples of phenotypic overlap caused by such biochemical convergence have yet to be uncovered in other types of MD, which are notorious for their clinically and biochemically heterogeneity^2^.

Beyond cytochrome *c*, we identify additional mitochondrial proteins with altered solubility that may contribute to the aforementioned mitochondrial dysfunctions. These include ABAT^84^, YRDC^85^, RMND1^86^, and MRPS16^87^ — factors involved in mtDNA stability, replication, and gene expression and mutated in genetic diseases. Reduced levels of mtDNA replication and maintenance proteins coincide with reduced mtDNA content measured at the onset of cardiomyopathy in males (14 weeks). Neither of these defects are rescued by blunting the mtISR via OMA1 inactivation in *Chchd10^S55L/+^Oma1^E324Q/E324Q^*mice(Figure 3i, j), suggesting that mtISR induction is not the primary driver of altered mtDNA homeostasis at the early stages of cardiac dysfunction. Not all identified proteins that exhibit reduced solubility are classical IMS residents, and thus their altered solubility may reflect broader proteostasis stress across mitochondrial compartments, which could be the consequence of altered mitochondrial ultrastructure and morphology^88^ (Figure 3g, h). CHCHD10^S55L^ aggregation may exert a ripple effect across mitochondrial homeostasis leading to the proteomic remodeling characterized by reduced levels of most proteins in isolated cardiac mitochondria. Alternatively, proteomic remodeling may directly contribute to mitochondrial membrane remodeling via the impaired assembly or maintenance of the MICOS complex (Figure 2k, m), which previously implicated in CHCHD10^S59L^-mediated cristae dysfunction^89,90^. Indeed, future studies are needed to dissect the individual contributions of mtDNA depletion, cristae dysmorphology, and impaired cytochrome *c* oxidation for cardiac function. This will be difficult given the interdependent nature of these mitochondrial processes^91^, yet emerging chemical approaches that target specific mitochondrial dysfunctions may prove to be useful in this endeavor ^89,92–94^.

At the outset, *Chchd10* mutant mice were developed to study the impact of the S59L pathogenic variant in *CHCHD10* patients suffering from autosomal dominant ALS and Frontotemporal Dementia (FTD)^25,26^. While there is little clinical evidence to support the existence of cardiac defects in the few *CHCHD10^S59L/+^* patients reported thus far, it is possible that subclinical, underlying molecular defects similar to those we and others reported in presymptomatic *Chchd10^S55L/+^* mutant cardiac mitochondria have yet to be discovered. Indeed, several MD nuclear genes (e.g. *SCO2, ANT1, SUCLA2, COX10, COX15*) that are mutated in patients with neurological abnormalities can also lead to cardiomyopathy^50,95–103^, underscoring the broader and evolving spectrum of clinical manifestations associated with MD. In *Chchd10^S55L/+^* mice, our findings confirm several previously described mitochondrial and cardiac dysfunctions caused by the toxic gain-of-function CHCHD10^S55L^ protein. However, several discrepancies with previous studies stand out, which could be explained by differences in mouse genetic backgrounds on which mutant mice were generated. In our study, we used heterozygous *Chchd10^S55L/+^* mice maintained on a C57BL/6N (N) background, whereas studies from the Manfredi and Narendra labs used the C57BL/6J (J) strain^21,25,28^. Genetic variations between these two sub-strains of C57Bl/6 mice are known to influence mitochondrial metabolism and cardiac health^104,105^. On the J-strain, cardiomyopathy onset occurs around 18 weeks in male *Chchd10^S55L/+^*mice, with mtDNA depletion and OXPHOS defects reported at one year^102^. In line, double deletion of *Chchd10* and *Chchd2*, which phenocopies mitochondrial defects observed in *Chchd10^S55L/+^*mice, causes cardiac dysfunction on the J-strain with kinetics similar to that of *Chchd10^S55L/+^* mice maintained on the same genetic background^106^. By contrast, cardiomyopathy manifests in *Chchd10^S55L/+^* mice on the N strain one month earlier^26^ (Figure 1b, c) concomitant with the depletion of mtDNA and impaired respiration. J-strain *Chchd10^S55L/+^* mice exhibit iron deficiencies^21^ while N-strain *Chchd10^S55L/+^* mice do not (Figure S2g), although it should be noted that the techniques used to measured biological metals (including iron) were different. Another divergence from prior studies is the emergence of sex-specific phenotypes in our model. Cardiac phenotyping studies on the J strain focused exclusively on females^25^ or males^28^, making it challenging to assess the influence of biological sex. However, in the single study of *Chchd10^S55L/+^* mice that included both sexes, Genin and coworkers reported that female mutants had left ventricular ejection fraction (%LVEF) values similar to wild-type male littermates at 23 weeks^26^. In line, we observe cardiac protection in *Chchd10* females (Figure 1b, c) and sexually dimorphic effects in *Chchd10^S55L/+^* mice on the N strain including a male-specific increase in cardiac inflammatory pathways. Environmental context may also contribute to differences observed between groups studying *Chchd10^S55L/+^* mice. Animal housing conditions, diet, and microbiota composition are increasingly recognized as critical modifiers of phenotype onset and severity^107^, which is likely relevant for mitochondrial disorders^108^. Recent work from the Manfredi group identified high fat diet feeding as a nutritional intervention capable of alleviating mtISR and cardiomyopathy in male mutant male mice by restoring CHCHD10 solubility to the aggregation-prone S55L variant^39,105,106^, which can also suppress cardiomyopathy in cardiomyocyte-specific *Yme1l1* knockout mice (*Yme1l1^Heart^*)^39,106,107^. These observations underscore the need to consider both genetic and non-genetic modifiers in interpreting preclinical disease models.

Finally, functional suppression of STING and OMA1 in *Chchd10^S55L/+^* mutant mice have provided insights into the relevance of overactive innate immune signaling and mtISR, respectively. The beneficial effects of STING ablation we observe likely reflect the general protection against overactive innate immune signaling triggered by mitochondrial dysfunction, which has been previously documented in mouse models for *Polg*, *Tfam*, *Pink1*, *Parkin*^39,109,110^ and in mice subjected to chemically-induced mitochondrial dysfunction^42,111^. For OMA1, our findings refine the understanding of the physiological relevance of the stress-induced metalloprotease in CHCHD10 pathogenesis. In the hearts of *Chchd10^S55L/+^*mice, OMA1 inactivation transiently preserves cardiac function without ultimately rescuing OXPHOS defects, mtDNA levels, or CHCHD10 protein solubility. This suggests that ISR suppression alone is insufficient to correct the primary energetic defect in the heart and instead supports a paradigm in which CHCHD10 aggregation and proteostatic dysfunction are the proximal triggers of mitochondrial dysfunction in the heart. Transcriptomic analyses reveal that OMA1 inhibition does not suppress overactive innate immune signaling in *Chchd10^S55L/+^ Oma1^E324Q/E324Q^* mutant hearts (Supplemental Dataset 6), which is consistent with the breadth of cardioprotection we observe. On the other hand, in skeletal muscle of mutant mice modeling the *CHCHD10^G58R^* allele, OMA1 deletion is synthetically lethal and exacerbates myopathy^106^.

We posit that these discrepancies stem from the nature of the toxic, gain-of-function CHCHD10 mutation (G58R versus S59L) and the affected tissue (skeletal muscle versus heart), rather than the difference between the deletion of *Oma1* and the catalytic inactivation by the *Oma1^E324Q/E324Q^*model, since the *Oma1^-/-^* and *Oma1^E324Q/E324Q^*cardiac proteomes are no different (Figure S3d)^108^. Indeed, future studies are needed to explore the breadth of protection offered by the *Oma1^E324Q/E324Q^*model, including in the neuromuscular system of *Chchd10^S55L/+^*mice and as well as other MD mouse models of intramitochondrial protein aggregation and proteostatic imbalance.

## Materials and Methods

### Animals

Animal care was conducted in accordance with European animal welfare laws (Directive 2010/63/EU). The French Ministry of Research and local Animal Ethics Committees reviewed and approved all experiments under the authorized protocol APAFIS #32988-2021091417302194. Mice were kept in a specific pathogen-free environment with standard conditions, including a 14-hour light/10-hour dark cycle, 50-70% humidity, and a temperature range of 19-21°C. They had unlimited access to food and water in cages equipped with bedding and gnawing sticks for enrichment. The principles of the 3Rs were followed to meet animal welfare standards. Animals were checked weekly, and euthanasia was performed to minimize pain and distress if body weight loss exceeded 20%. The generation of *Chchd10^S55L/+^*mice was described previously^109^. *Oma1^E324Q/E324Q^* mice were generated using CRISPR/Cas9 endonuclease-mediated genome editing on the C57Bl6/N background. The sgRNA 5’-GTGCGATCTCATGGCCCAGG *AGG*-3’ was designed using the CRISPOR Web tool (http://crispor.tefor.net/)^112^.

The donor matrix chosen was a 161 nucleotide ssODN: (GAATGGACAAGTGTTTATTTTCACCGGGCTTCTGAATAGTGTGACGGACGTGCACCACTGTCCTTtCTCCTGGGCCATcAGATCGCACACGCAGTCCTGGGGCACGCCGTGAGTACCGGGATGTGCAGCTGCTGAATGTTTGCTTTGATATTAGCAAGTG).

For transgenesis experiments, 3-week-old C57BL/6N females were superovulated and mated with C57BL/6N stud males to recover one-cell embryos. Pronuclei of fertilized embryos were co-microinjected with RNP CRISPR/Cas9 complex (20 ng/μL Cas9 protein, 20 ng/ μL sgRNA) together with 20 ng/μl of ssODN (all reagents from Integrated DNA Technologies, IDT). Microinjected embryos were then implanted into the oviducts of C57BL/6J×CBA F1 foster mothers following standard procedures^112^.

F0 Mutant mice were screened by PCR and confirmed by direct sequencing^113^ followed by germline transmission analysis and confirmation of the descendants for mutational insertions and off target effects. *Chchd10^S55L^ Oma1^E324Q/E324Q^* were backcrossed on C57Bl6/N background. *Sting^Gt/Gt^* mice (IMSR_JAX:017537) and *Chchd10^S55L^ Sting^Gt/Gt^* mice were intercrossed backcrossed on C57Bl6/N background. Cardiomyocyte-specific *Yme1l1* knockout mice (*Yme1l^Heart^*)^110^ were rederived on a C57Bl6/N background.

### Echocardiography

Transthoracic echocardiography was conducted using a Vevo 3100 Imaging System paired with a 25-55 MHz linear-frequency transducer (MX550D, FUJIFILM VisualSonics). Randomized WT, *Chchd10^S55L/+^*, *Oma1^E324Q/E324Q^*, *Chchd10^S55L/+^Oma1^E324Q/E324Q^*, *Sting^Gt/Gt^* and *Chchd10^S55L/+^Sting^Gt/Gt^* mice were anesthetized with 2% isoflurane in oxygen and positioned supine on a 37°C heated pad. Limb electrodes and a rectal probe monitored the ECG and body temperature. Prior to echocardiography and applying ultrasound gel, the fur on the thorax was removed with hair-removal cream. To evaluate left ventricle (LV) size and function, B– and M-Mode images were captured in the parasternal long-axis view (PLAX) at a heart rate of 400-500 bpm. Measurements of the systolic and diastolic LV dimensions, including interventricular septum thickness (IVS; mm), LV diameter (LVD; mm), LV posterior wall thickness (LVPW; mm), and cardiac output [ejection fraction (% LVEF)], were obtained by analyzing at least three independent cardiac cycles across at least three M-Mode images using Vevo Lab (VisualSonics), as previously described^73^.

### Histology

Mice were euthanized through cervical dislocation, and their hearts were removed post-mortem. The entire hearts were fixed in 4% formaldehyde (VWR chemicals) overnight and then fully dehydrated using a series of ethanol gradients. The tissues were then embedded in paraffin and sectioned into 4 μm thick slices using a microtome. These sections were deparaffinized in xylene, rehydrated, and subsequently stained with hematoxylin and eosin (H&E) or Picrosirius Red (ABCAM, ab150681) according to standard protocols. The images were captured using a slide scanner AxioScan.Z1 (ZEISS). Histopathological evaluation was performed using digitized whole-slide images and the QuPath software^114^. Images were analyzed using Fiji (ImageJ) by quantifying the area in red as fibrosis area and normalized by the total area of the tissue section using a macro. All analysis settings and annotations were stored within the same project. The annotation tool was used to segment the tissue area for each image. To determine the extent of fibrosis, we used QuPath’s machine learning-based object classifier. Annotation classes were defined for both fibrosis and background areas. For model training, three representative cropped regions were selected from each image included in the analysis, and a composite training set was created. Hundreds of annotations were added for each class to ensure robust classifier performance. Particles smaller than 2 µm were excluded from the fibrosis area analysis. Finally, the ratio of the fibrosis area to the total tissue area was calculated and represented.

### SDS-PAGE

Immunoblot analysis was conducted to evaluate the steady-state protein levels in cardiac tissue. For tissue lysates, mice were euthanized via cervical dislocation, followed by opening the chest, excising the hearts, weighing them, flash-freezing them in liquid nitrogen, and storing them at 80°C. Tissue were prepared by homogenizing them in cold RIPA buffer [1 mg/20 µL, 1% Triton X-100, 1% sodium deoxycholate, 0.1% SDS, 150 mM NaCl, 50 mM Tris·HCl (pH 7.8), 1 mM EDTA, and 1 mM EGTA] with protease and phosphatase inhibitors, and kept on ice for 30 minutes. The homogenate was then centrifuged for 15 minutes at 16,000g, 4°C. Protein concentration was measured by the Bradford assay (Bio-Rad) using a BSA standard curve, with absorbance read at 595 nm using an Infinite M2000 microplate reader (Tecan). Equal amounts of protein were mixed with 4x Laemmli Sample Buffer [355 mM, 2-mercaptoethanol, 62.5 mM Tris-HCl pH 6.8, 10% (v/v) glycerol, 1% (w/v) SDS, 0.005% (v/v) Bromophenol Blue], then heated at 70°C for 5 minutes. Samples (10 µg) were separated on 4-20% polyacrylamide gels (Mini Protean TGX Stain-Free gels, BioRad) and transferred to nitrocellulose membranes using the Trans-Blot Turbo Transfer system (Bio-Rad). Consistent protein loading across lanes was verified using Ponceau S staining or Stain-free detection. Membranes were blocked for 2 hours with 5% (w/v) semi-skimmed dry milk in Tris-buffered saline with 0.1% Tween (TBST), then incubated overnight at 4°C with primary antibodies (Supplemental Dataset 4) diluted 1:1,000 in 2% (w/v) Bovine Serum Albumin (BSA) in 0.1% TBST. The following day, membranes were incubated with HRP-conjugated secondary antibodies at room temperature for 2 hours (diluted 1:10,000 in 2% BSA TBST 0.1%). Finally, membranes were treated with Clarity Western ECL Substrate (Bio-Rad) for 2 minutes, and luminescence was detected using the ChemiDoc Gel Imaging System. Densitometric analysis of the immunoblots was performed using Image Lab Software v.6.1.0 (Bio-Rad).

### Mitochondrial isolation

Freshly isolated cardiac mitochondria were prepared as previously detailed^73^. In summary, ventricles were separated from atria and non-cardiac tissues, chopped into small pieces, and then manually homogenized in an ice-cold 2-ml homogenizer with IB buffer (275 mM sucrose, 20 mM Tris, 1 mM EGTA-KOH, pH 7.2) containing Trypsin-EDTA (0.05%). To inhibit trypsin activity, bovine serum albumin (BSA) fatty acid-free (0.25 mg/mL) and protease inhibitor cocktail (PIC, Roche) were added. The cardiac homogenates were first centrifuged at low speed (1000 g, 10 min, 4 °C) to eliminate nuclei and debris, then centrifuged again at a higher speed (3200 g, 15 min, 4 °C) to isolate the crude mitochondrial fraction. The crude mitochondrial pellet was resuspended in IB buffer, and the protein concentration was measured using the Bradford assay.

### High resolution (Fluo)respirometry

Oxygen consumption and mitochondrial membrane potential were measured in cardiac mitochondria using the High-Resolution Respirometry system (O2k-Fluorespirometer, Oroboros, AT). For this, freshly isolated cardiac mitochondria from adult WT, Chchd10*^S55L^*, *Oma1^E324Q/E324Q^*and Chchd10*^S55L^/Oma1^E324Q/E324Q^* mice (7 or 14 weeks of age) were used, as described above. The respiration and membrane potential (Δψ) of the mitochondria were analyzed using O2K-Fluorometry with the O2K-Fluorescence LED2-Module connected via the amperometric channel of the O2K. Briefly, 50 µg of cardiac mitochondria were suspended in Mir05 buffer [MgCl_2_-6H_2_O 3 mM, Lactobionic Acid 60 mM, Taurine 20 mM, KH_2_PO_4_ 10 mM, Hepes-KOH 20 mM, Sucrose 110 mM, EGTA-KOH 0.5 mM, BSA (1g/L)]. Rhodamine 123 (RH-123) (0.66 µM) fluorescence quenching (Δ fluorescence) was used to measure the membrane potential (Δψ) in energized mitochondria. Maximal mitochondrial respiration capacity (OXPHOS) was determined by adding PGM [10 mM pyruvate, 5 mM glutamate, 5 mM malate, state 2] or malate (2 mM) and palmitoyl-carnitine (10 µM) in the presence of ADP (1 mM, state 3) to measure complex I-driven respiration; rotenone (0.5 µM) and succinate (10 mM, state 2) to measure complex II-driven respiration; and antimycin A (2.5 µM), ascorbate (2 mM), *N,N,N′,N′-tetramethyl-p-phenylenediamine* (TMPD, 0.5 mM), and carbonyl cyanide m-chlorophenyl hydrazone (CCCP, 2 µM) to measure complex IV-driven respiration. Oxygen consumption was assessed using 50 µg of mitoplasts – cardiac mitochondria subjected to a freeze–thaw cycle – resuspended in Mir05 buffer, following the sequential addition of antimycin A (2.5 µM), ascorbate (2 mM), *N,N,N′,N′-tetramethyl-p-phenylenediamine* (TMPD, 0.5 mM), and bovine cytochrome c (10 µM). The results were analyzed using Datlab v.7.

### Alkaline Carbonate Extraction

Alkaline carbonate extraction of membrane proteins was carried out as previously reported^115^. Crude mitochondria, isolated from mouse hearts, were resuspended and treated with 0.1 M sodium carbonate (Na₂CO₃) at pH levels of 12.5 or 9.5, and incubated on ice for 30 minutes. The mixtures were then ultracentrifuged at 45,000 rpm for 30 minutes at 4°C using Beckman polycarbonate tubes and a TLA 55 rotor. Both the supernatants and pellets were treated with a trichloroacetic acid buffer (17.5% TCA, 40 mM HEPES, 0.02% Triton X-100) on ice for 30 minutes, followed by centrifugation at 21,130g for 20 minutes at 4°C. The resulting samples were washed three times with cold 100% acetone and left to air-dry at room temperature for 30 minutes. Finally, the dried pellets were resuspended in 1× Laemmli sample buffer (Bio-Rad) for analysis via SDS-PAGE and western blot.

### Differential solubility proteomics

To obtain soluble and insoluble mitochondrial protein fractions, 150 µg of mitochondria were pelleted at 20,000 g for 5 min at 4°C in two separate tubes. One pellet was resuspended in 45 µL 2X Laemmli buffer (total fraction). The second pellet was resuspended in 37.5 µL Fractionation Buffer A (10 mM TRIS-HCl, pH=8; 1 mM EDTA; 1% Triton X-100 (1% v/v); cOmplete protease inhibitor (Roche) and incubated at 4°C for 10 min. After centrifugation at 20,000 g for 10 min at 4°C. 37.5 µL of the supernatant was transferred to a fresh tube (soluble fraction) and supplemented with 7.5 µL 4X Laemmli buffer. The insoluble pellet was washed twice with 150 µL Fractionation Buffer A and subsequently centrifuged at 20,000 g for 10 min at 4°C. The supernatant was discarded, and the pellet was resuspended in 45 µL Urea sample buffer (125 mM TRIS-HCl, pH=6.8; 6M Urea, 6%SDS; 0.5% beta-mercaptoethanol; 0.01% bromphenol blue). The individual fractions total (mitochondria), pellet and supernatant were subjected for protein digestion and LC-MS/MS measurement.

### Protein Digestion

One third of each fraction (total, pellet, supernatant) were subjected for trypsin digestion. In detail, proteins were reduced (10 mM TCEP) and alkylated (20 mM CAA) in the dark for 45 min at 45 °C. Samples were subjected to an SP3-based digestion^116^. Washed SP3 beads (SP3 beads (Sera-Mag(TM) Magnetic Carboxylate Modified Particles (Hydrophobic, GE44152105050250), Sera-Mag(TM) Magnetic Carboxylate Modified Particles (Hydrophilic, GE24152105050250) from Sigma Aldrich) were mixed equally, and 3 µL of bead slurry were added to each sample. Acetonitrile was added to a final concentration of 50% and washed twice using 70 % ethanol (V=200 µL) on an in-house manufactured magnet. After an additional acetonitrile wash (V=200µL), 5 µL digestion solution (10 mM HEPES pH = 8.5 containing 0.5µg Trypsin (Sigma) and 0.5µg LysC (Wako)) was added to each sample and incubated overnight at 37°C. Peptides were desalted on a magnet using 2 x 200 µL acetonitrile. Peptides were eluted in 10 µL 5% DMSO in LC-MS water (Sigma Aldrich) in an ultrasonic bath for 10 min. Eluted tryptic peptides subjected to further peptide purification using the StageTip technique using the SDB material. Samples were stored at –20°C and 10 µL of 2.5% formic acid and 2% acetonitrile were added. 3 µL were used for a LC-MS/MS run.

### Liquid Chromatography and Mass Spectrometry

LC-MS/MS instrumentation consisted of an Easy-LC 1200 (Thermo Fisher Scientific) coupled via a nano-electrospray ionization source to an Exploris 480 mass spectrometer (Thermo Fisher Scientific, Bremen, Germany). An Aurora Frontier column (60 cm length, 1.7 µm particle diameter, 75 µm inner diameter, Ionopticks). A binary buffer system (A: 0.1 % formic acid and B: 0.1 % formic acid in 80% acetonitrile) based gradient was utilized as follows at a flow rate of 185 nL/min; a linear increase of buffer B from 4% to 28% within 100 min, followed by a linear increase to 40% within 10 min. The buffer B content was further ramped to 50 % within 4 minutes and then to 65 % within 3 minutes. 95 % buffer B was kept for a further 3 min to wash the column. The RF Lens amplitude was set to 45%, the capillary temperature was 275°C and the polarity was set to positive. MS1 profile spectra were acquired using a resolution of 30,000 (at 200 m/z) at a mass range of 450-850 m/z and an AGC target of 1 × 106. For MS/MS independent spectra acquisition, 34 equally spaced windows were acquired at an isolation m/z range of 7 Th, and the isolation windows overlapped by 1 Th. The fixed first mass was 200 m/z. The isolation center range covered a mass range of 500–740 m/z. Fragmentation spectra were acquired at a resolution of 30,000 at 200 m/z using a maximal injection time setting of ‘auto’ and stepped normalized collision energies (NCE) of 24, 28, and 30. The default charge state was set to 3. The AGC target was set to 3e6 (900% – Exploris 480). MS2 spectra were acquired in centroid mode. FAIMS was enabled using an inner electrode temperature of 100°C and an outer electrode temperature of 90°C. The compensation voltage was set to –45V.

Raw files were analyzed using Spectronaut 19.3.241023.62635 in direct DIA mode using the Uniprot Mus musculus (Mouse) UP000005640 proteome – one fasta sequence per gene, reviewed, 21,984 protein sequences). Trypsin/P was selected as the cleavage rule using a specific digest type. The minimal peptide length was set to 7 and a total of 2 missed cleavages were allowed. The peptide spectrum match (PSM), peptide, and protein group FDR were controlled to 0.01. The mass tolerances were used with default settings (Dynamic,1). The directDIA+(deep) workflow was selected and cross-run normalization (only for the whole proteome analysis) was enabled.

For whole proteome analysis (total fraction), the protein group file was exported and LFQ intensities (MaxLFQ algorithm)^117^ were log2 transformed. Statistically significant different proteins were identified using a two-sided t-test followed by a permutation-based FDR calculation (s0=0.1, number permutations=500, FDR< 0.05) using Instant Clue^118^.

ISR genes were annotated according to Class 1 and ATF-dependent upregulated genes defined previously^116^.

For solubility proteome analysis, the iBAQ (intensity based absolute quantification) intensity was used. The individual fractions (total, pellet, supernatant) were median normalized within the groups. Then the pellet fraction was calculated as iBAQ total / iBAQ pellet.

### Inductively coupled plasma optical emission spectroscopy

Heart tissue samples (9–30 mg) and mitochondrial preparations (equivalent to 100 µg protein) were analyzed for elemental content using inductively coupled plasma–optical emission spectrometry (ICP-OES; Optima 7300DV, PerkinElmer Life Sciences). Samples were digested in 40% nitric acid by boiling for 1 hour in acid-washed, semi-sealed tubes. After digestion, samples were diluted with ultrapure, metal-free water (18.2 MΩ·cm) prior to analysis. Emission intensities for Ca, Cu, Fe, K, Mg, Mn, P, S, and Zn were measured simultaneously, with acid matrix blanks prepared identically to the samples for background correction. Each biological replicate (n = 5–6 per genotype and sex) was measured in technical triplicate, and the average intensity calculated by area under the curve was used for quantification. Elemental concentrations were calculated using three-point standard curves generated from serial dilutions of two commercially available mixed-metal standards (Optima). Blanks of the nitric acid matrix, with and without metal spikes, were included to confirm the reproducibility and accuracy of measurements.

### Heme quantification by HPLC

Mitochondria equivalent to 100 µg of protein were used for heme analysis by HPLC. Heme was extracted as previously described^119^. Mitochondrial pellets were resuspended in 100 µL of 97.5% acetone / 2.5% hydrochloric acid (v/v), vortexed for 1 minute, and centrifuged at 12,000 × g for 5 minutes at room temperature. The supernatant was carefully transferred to a new tube to avoid disturbing the pellet. This acid-acetone extract was mixed with 100 µL acetonitrile, 1 µL formic acid, and 7 µL ammonium hydroxide, then vortexed briefly and centrifuged again at 12,000 × g for 2 minutes. An aliquot of 100 µL of the final solution was injected onto a Sonoma C18 reversed-phase HPLC column (C18(2) 10µ 100Å 25 cm x 4.6 mm), equilibrated in 0.05% trifluoroacetic acid with 30% acetonitrile. Chromatographic separation was achieved using a 5-minute linear gradient from 30% to 50% acetonitrile, followed by a 20-minute gradient from 50% to 75%, at a flow rate of 1 mL/min. The column was subsequently washed with 99% acetonitrile 0.05% trifluoroacetic acid then re-equilibrated with starting buffer. Heme elution was monitored at 405 nm via the Soret band. Retention times were determined using purified standards of heme b, heme o, and heme a. Quantification was based on the area under the curve, with only heme b concentrations reported.

### Cardiac RT-qPCR and bulk RNAseq

Total RNA was isolated from snap-frozen left ventricles by the NucleoSpin RNA kit (Macherey-Nagel, 740955). Quality control was performed on an Agilent BioAnalyzer. Libraries were built using a TruSeq Stranded mRNA library Preparation Kit (Illumina, USA) following the manufacturer’s protocol. For the Control (*Yme1l1^LoxP/LoxP^*) and Y*me1l1^Heart^* mice samples, two runs of RNA sequencing were performed for each library on an Illumina NextSeq 500 platform using single-end 75bp. For the *Chchd10^S55L/+^* and *Oma1^E324Q/E324Q^*mutant mice, a single run was performed for all libraries on an Illumina NextSeq 2000 platform using paired-end 52b reads. The RNA-seq analysis was performed with Sequana 0.18.1. In particular, we used the RNA-seq pipeline (version 0.20.0) (https://github.com/sequana/sequana_rnaseq) built on top of Snakemake 7.32.4^119^. Reads were trimmed from adapters using Fastp 0.23.2 then mapped to the mouse reference genome GRCm39 using STAR 2.7.10a^120^. FeatureCounts 2.0.1 was used to produce the count matrix, assigning reads to features using annotation from Ensembl GRCm39_113 with strand-specificity information^121^. Quality control statistics were summarized using MultiQC 1.16.0^122^. Clustering of transcriptomic profiles were assessed using a Principal Component Analysis (PCA). Differential expression testing was conducted using DESeq2 library 1.38.3.0^123^ scripts indicating the significance (Benjamini-Hochberg adjusted p-values, false discovery rate FDR < 0.05) and the effect size (fold-change) for each comparison. Over-representation analysis (ORA) was performed to determine if genes modulated by genotype or biological sex are more present in specific pathways. ORA was performed on WebGestalt (https://www.webgestalt.org/). RNAseq data have been deposited at ENA with the dataset identifier E-MTAB-15304 and E-MTAB-15305. For RT-qPCR, 1 µg of total RNA was converted into cDNA using the iScript Reverse Transcription Supermix (Bio-Rad). RT-qPCR was performed using the CFX384 Touch Real-Time PCR Detection System (Bio-Rad) and SYBR® Green Master Mix (Bio-Rad) using the primers listed in Supplemental Dataset 8. *Gapdh* was amplified as internal standard. Data were analyzed according to the 2−ΔΔCT method^124^.

### mtDNA quantification

Genomic DNA was isolated using the NucleoSpin Tissue kit (MACHEREY-NAGEL) and quantified with a NanoQuant Plate (Infinite M200, TECAN). Quantitative PCR (qPCR) was carried out using the Real-Time PCR Detection System (Applied Biosystems StepOnePlus), with 20 ng of total DNA and SYBR Green Master Mix (Bio-Rad). β-*Actin* was amplified as an internal nuclear gene control, as previously described^125^, using primers listed in Supplemental Dataset8. Data were analyzed according to the 2^−ΔΔCT method^126^.

### Transmission Electron Microscopy

Transmission electron microscopy was conducted on cardiac tissue mice at 14 weeks of age, as previously described^73^. Small tissue samples (1 x 1 x 1 mm) from the posterior wall of the left ventricle were fixed initially in a 37 °C pre-warmed solution containing 1x PHEM buffer (60 mM PIPES, 25 mM HEPES, 10 mM EGTA, 2 mM MgCl₂, pH 7.3), 2.5% glutaraldehyde, and 2% paraformaldehyde (PFA) for 30 minutes, followed by overnight fixation at 4 °C. The specimens were rinsed three times with 3x PHEM buffer, and then were processed following an adapted OTO protocol^127^. First, samples were incubated in a mixture of 1% osmium tetroxide (OsO₄) and 1.5% potassium ferrocyanide (K₄Fe(CN)₆) to enhance membrane contrast. Then, samples were treated with 1% tannic acid to further increase contrast. A second post-fixation step was performed with 1% OsO₄ alone. Samples were dehydrated through a graded ethanol series and embedded in SPURR resin (Electron Microscopy Sciences) at 60°C for 48 hours. Thin sections (70 nm) were then cut with a Leica UCT microtome and collected on carbon and formvar-coated copper grids. The sections were contrasted with 4% aqueous uranyl acetate and Reynold’s lead citrate. The images were acquired using a TECNAI T12 Transmission Electron Microscope (FEI), operated at 120 kV with a RIO16 camera (Gatan), controlled by Digital Micrograph software. Mitochondrial ultrastructure quantification was performed using NIH ImageJ software, as previously described^128^. Mitochondrial area and number were quantified from electron micrographs, with 315–440 mitochondria analyzed per experimental group. The same set of mitochondria was used to assess cristae morphology using a standardized scoring system, as described previously^129^.

### 1D BN-PAGE of cardiac mitochondria

One-dimensional blue native PAGE (1D BN-PAGE) was performed following a previously described^130^ with some modifications. In summary, heart mitochondria (50 μg, with protein concentration determined using the DC Protein Assay from BIO-RAD) were isolated from WT, *Chchd10^S55L^*, *Oma1^E324Q/E324Q^*, *Chchd10^S55L^Oma1^E324Q/E324Q^*mice. These mitochondria were incubated with a digitonin extraction buffer composed of HEPES (30 mM), potassium acetate (150 mM), glycerol (12%), 6-aminocaproic acid (2 mM), EDTA (1 mM), and high-purity digitonin (10%), with a pH of 7.2. The mitochondria were vortexed for 1 hour at 4°C to solubilize the membranes, followed by centrifugation at 21,130g for 30 minutes. The supernatant was collected and mixed with loading dye containing Coomassie Brilliant Blue G-250 (0.0125% w/v, InvitrogenTM, BN2004). The solubilized mitochondria were loaded onto a 4-16% Bis-Tris acrylamide gel (1 mm, InvitrogenTM NovexTM NativePAGETM, BN2111BX10), using an anode buffer (Invitrogen, BN2001) with Cathode Buffer Additive (0.5%) mixed into the anode buffer (Invitrogen, BN2002). Electrophoresis was carried out at 80V and 20mA for 45 minutes, then at 150V and 20mA for 13 hours. Afterward, the gel was incubated in transfer buffer (0.304% w/v Tris, 1.44% w/v glycine) containing 0.2% SDS and 0.2% β-mercaptoethanol for 30 minutes at room temperature to denature the proteins. Following this, proteins were transferred to a polyvinylidene difluoride (PVDF) membrane in transfer buffer (0.304% w/v Tris, 1.44% w/v glycine, and 10% v/v ethanol) at 400mA and 20V for 3 hours and 30 minutes. The membrane was washed with methanol to remove the Coomassie stain. For immunodetection, the membrane was blocked with 5% milk in TBST (Tween-Tris-buffered saline) for 2 hours at room temperature, then incubated overnight with a specific primary antibody diluted in the blocking solution. The next day, the membrane was washed three times in TBST, then incubated with an HRP-conjugated secondary antibody for 2 hours at room temperature. Finally, the membrane was treated with Tris-HCl (0.1 M, pH 8.5) containing luminol and p-coumaric acid for 3 minutes, and luminescence was detected using a ChemiDoc TM XRS+ Imaging System. Band intensities were quantified using Image Lab Software.

## Statistical Analyses

Experiments were conducted at least three times, with quantitative analyses performed in a blinded manner. Group randomization (e.g., by genotype) was applied when simultaneous, parallel measurements were not feasible (e.g., Oroboros, cardiac isolation). For high-throughput assessments (e.g., proteomics, immunoblots, qRT-PCR), all groups were measured in parallel to minimize experimental bias. Statistical analyses were conducted using GraphPad Prism v10 software, and data are presented as mean ± SD or SEM, as specified. Statistical tests and replicate numbers are detailed in the figure legends. Comparisons between two groups were made using an unpaired two-tailed T-test, while one-way or two-way ANOVA was used to compare more than two groups or groups across multiple time points. Significance was set at P<0.05, with levels denoted as *P < 0.05, **P< 0.01, ***P< 0.001, and ****P< 0.0001.

## Data availability

All source data for the experiments are available with this manuscript. The datasets generated in this study have been deposited in the Proteomics Identification Database (PRIDE). Accession numbers are as follows: Differential solubility proteomics and proteome (PRIDE identifier: PXD064057 and PXD064045) and transcriptomics (ENA E-MTAB-15304 and E-MTAB-15305). The processed and analyzed data for proteomics included as Supplementary Datasets. Complete datasets for echocardiographic measurements analyzed in this study are not publicly available due to the limitations in exporting understandable file names from Vevo 3100 software (Visualsonics). However, these datasets are available upon request. Source data are included with this paper.

## Author contributions

M.A.C.R. performed in vivo phenotyping, histology sample preparation, RNA extraction, biochemical assays, mitochondrial respiration measurements, transmission electron microscopy (TEM) quantification, and data analyses. E.D. contributed to in vivo phenotyping, respiration assays, data analyses, mouse colony management, and coordination of animal facility logistics. H.N. conducted proteomic analyses, data processing, and data management. P.C. performed inductively coupled plasma (ICP) analyses and contributed to data interpretation. E.V. carried out qPCR and RT-qPCR, RNA extraction, data analyses, and participated in mouse colony management. D.M. prepared samples for proteomics analyses. J.D.H.C. developed image analysis methods and contributed to quantitative image data analyses. F.L.V. generated the *Oma1* mutant mouse model and coordinated animal facility operations. E.K. conducted RNA sequencing analyses and managed associated data. E.P. prepared TEM samples and acquired electron microscopy images. T.L. supervised proteomics experiments and contributed funding. V.P.F. provided the *Chchd10* mouse model and contributed funding. T.W. oversaw data analyses and management, secured funding, obtained GMO and animal experimentation permits, supervised the project, and wrote the manuscript, which was edited by all authors.

## Acknowledgments

We acknowledge David Hardy and coworkers at the Institut Pasteur Histology core and Stéphane Rigaud at the Image Analysis Hub for excellent service, as well as zootechnicians and veterinarians Jean Jaubert, Marion Berard, and Myriam Mattei at the Institut Pasteur Central Animal Facility. We thank Marie Lemesle for administrative assistance and Scot Leary, Alice Lepelley, Emmanuelle Genin, and all the members of the Wai lab for helpful discussions. MACR, TW and VPF are supported by the FRM (MitoDeath). JDHC is supported by the Institut Pasteur Roux-Cantarini Fellowship. TW is supported by the Agence Nationale de la Recherche (ANR-21-CE14 0052-02, ANR-23-CE13-0043-01), and Institut Pasteur (INNOV 164-2022). We are grateful for support for Ultrastructural BioImaging Core Facility equipment from the GIS-IBISA, the French Government Programme Investissements d’Avenir France BioImaging (FBI, N° ANR-10-INSB-04-01), the DIM1Health and the French gouvernement (Agence Nationale de la Recherche) Investissement d’Avenir programme, Laboratoire d’Excellence “Integrative Biology of Emerging Infectious Diseases” (ANR-10-LABX-62-IBEID).

## Supplemental Information

**Supplemental Figure 1** – Cardiac remodeling in *Chchd10* mutant hearts

**Supplemental Figure 2** – Impaired mitochondrial respiration in mutant *Chchd10* hearts

**Supplemental Figure 3** – OMA1 and mtISR activation

**Supplemental Figure 4** – Differential solubility proteomics in *Chchd10* mice

**Supplemental Dataset 1** – Bulk RNAseq

**Supplemental Dataset 2** – Pathway and Gene analyses Chchd10_vs_WT_and_M_vs_F.

**Supplemental Dataset 3** – Pathway Analyses RNAseq_Chchd10_vs_WT_Males

**Supplemental Dataset 4** – Pathway Analyses RNAseq_Chchd10_vs_WT_Females

**Supplemental Dataset 5** – Total Proteomics_4_Genotypes

**Supplemental Dataset 6** – Chchd10_Oma1_F_vs_Chchd10_F Pathway Analyses

**Supplemental Dataset 7** – Differential solubility cardiac proteomics

**Supplemental Dataset 8** – Reagents

