## Supplemental Information for "Mutant CHCHD10 disrupts cytochrome *c* oxidation and activates mitochondrial retrograde signaling in a model of cardiomyopathy"

**Title**

**Supplemental Information**

**Supplemental Figures 1-4**

Supplemental Dataset 1 – Bulk RNAseq

Supplemental Dataset 2 – Pathway and Gene analyses Chchd10_vs_WT_and_M_vs_F.

Supplemental Dataset 3 – Pathway Analyses RNAseq_Chchd10_vs_WT_Males

Supplemental Dataset 4 – Pathway Analyses RNAseq_Chchd10_vs_WT_Females

Supplemental Dataset 5 – Total Proteomics_4_Genotypes

Supplemental Dataset 6 – Chchd10_Oma1_F_vs_Chchd10_F Pathway Analyses

Supplemental Dataset 7 – Differential solubility cardiac proteomics

Supplemental Dataset 8 – Reagents

Raw Data

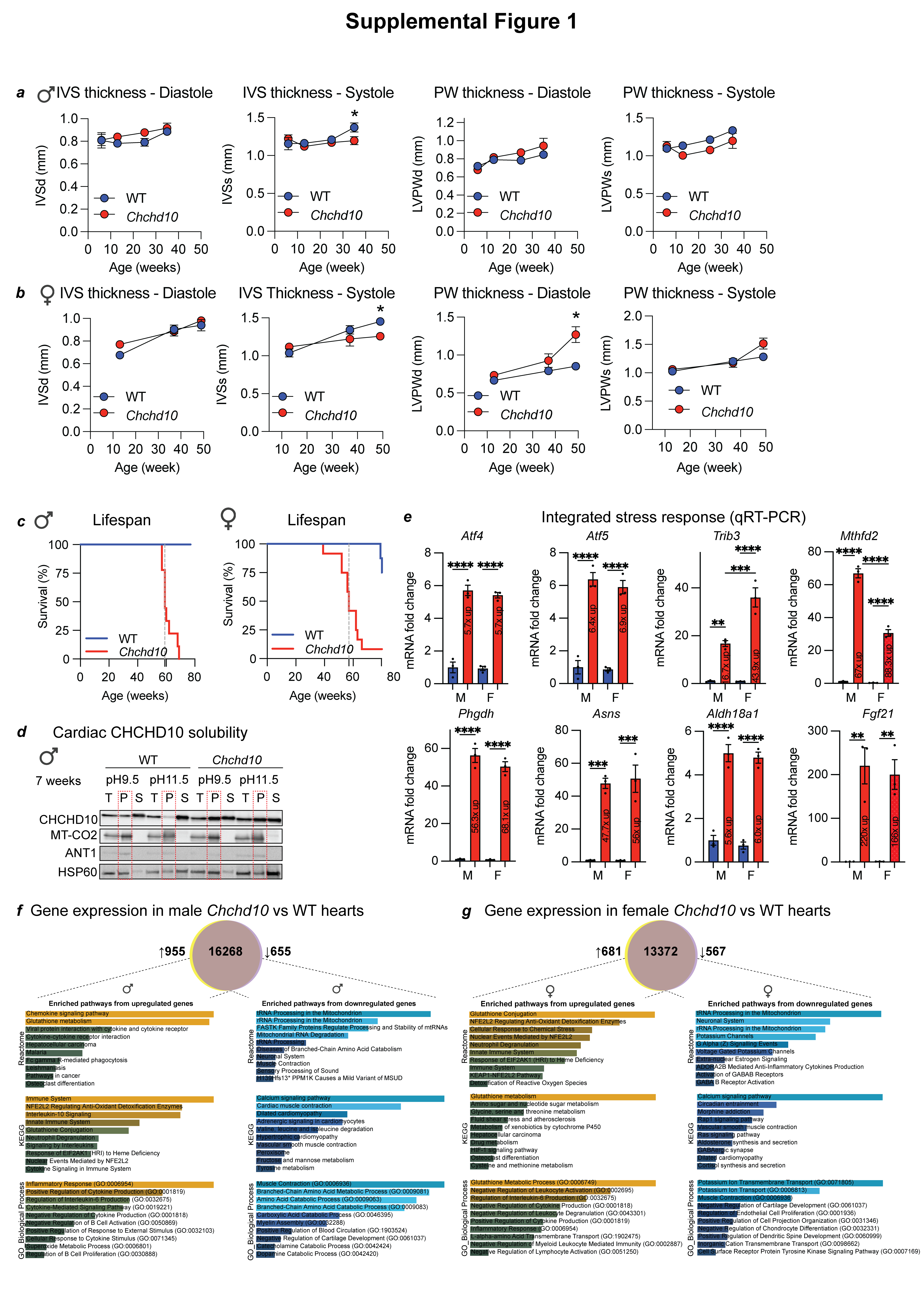

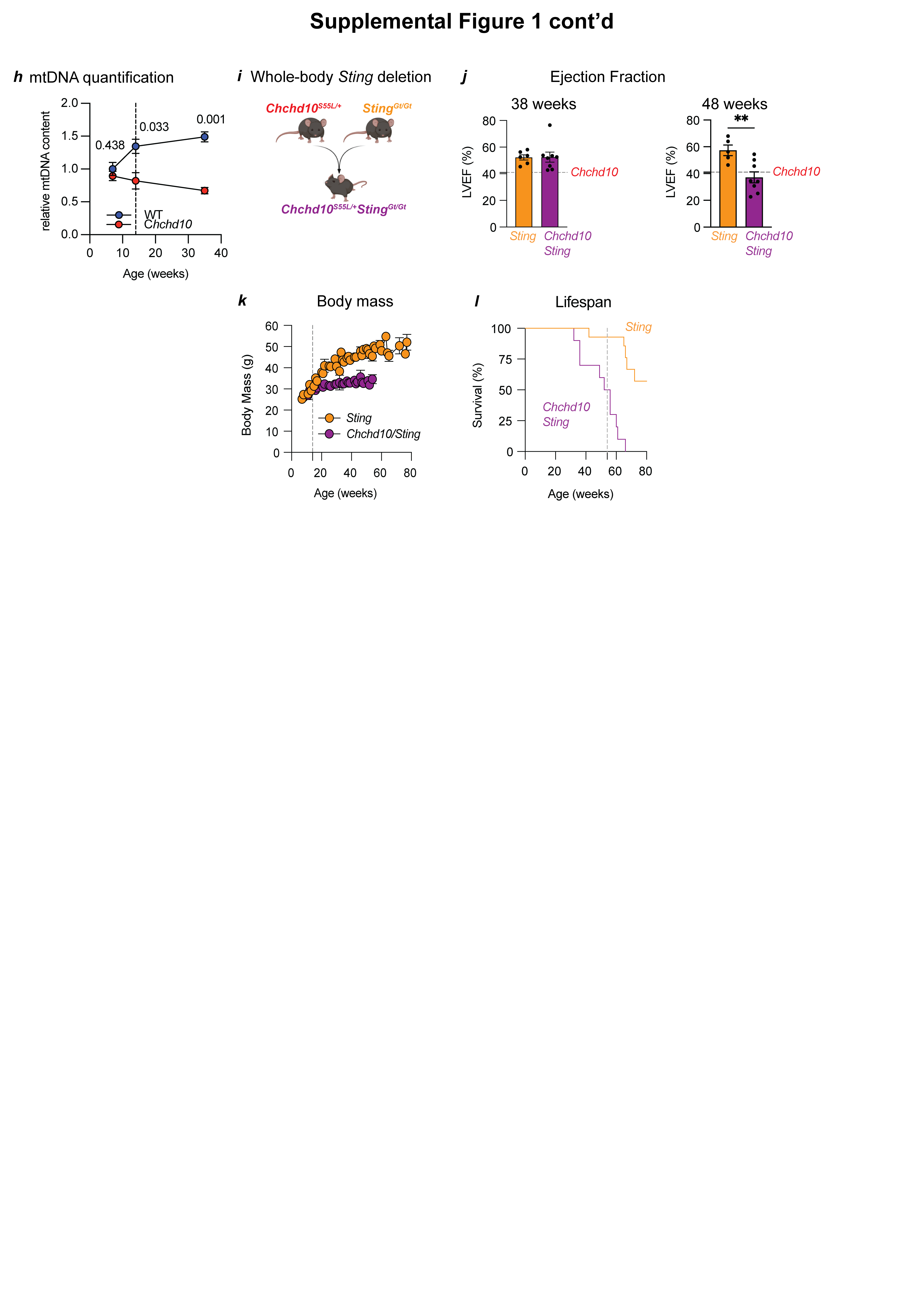
**Supplemental Figure 1 – Cardiac remodeling in *Chchd10* mutant hearts**

a) Diastolic (IVSd, mm) and Systolic interventricular septum thickness (IVSs, mm) and Diastolic (LVPWd, mm) and Systolic left ventricle posterior wall thickness (LVPWs, mm) of WT (blue, n=3-9) and *Chchd10* (red, n=3-8) male mice in Figure 1c. Data represent mean *±* SEM. One-way ANOVA, * P<0.05.
b) Diastolic (IVSd, mm) and Systolic interventricular septum thickness (IVSs, mm) and Diastolic (LVPWd, mm) and Systolic left ventricle posterior wall thickness (LVPWs, mm) of WT (blue, n=4-9) and *Chchd10^S55L^* (red, n=6-7) female mice in Figure 1c. Data represent mean *±* SEM. One-way ANOVA, * P<0.05.
c) Kaplan-Meier survival curve of WT (blue, n=10) and *Chchd10* (red, n=9) male mice (left) and WT (blue, n=13) and *Chchd10* (red, n=12) female mice (right). Dotted line (grey) represents median lifespan of *Chchd10* mice (male; 62 weeks, Log-rank test, P=0.0006 female 57 weeks, Log-rank test, P<0.0001).
d) Alkaline carbonate (Na_2_CO_3_) extraction of cardiac mitochondria performed on WT and *Chchd10* mutant male mice at 7 weeks of age. Total (T), insoluble pellet (P), and soluble supernatant (S) fractions were analyzed by immunoblotting with indicated antibodies.
e) Analysis of integrated stress response (ISR) genes *Atf4*, *Atf5*, *Trib3*, *Mthfd2*, *Phgdh*, *Asns, Aldh18a1,* and *Fgf21* via qRT-PCR of total cardiac biopsies from wild type (blue, n=3) and *Chchd10* (red, n=3) male and female mice at indicated ages. Data are relative mean fold changes ± SEM, 2-tailed unpaired Student’s t test. **P<0.01, ***P<0.001,****P<0.0001.
f) Venn diagram of differentially expressed genes (DEGs) in male *Chchd10*  versus WT mice. Bulk RNA-seq was performed on cardiac biopsies (n=3) at 14 weeks of age. 955 genes were upregulated in male *Chchd10* vs wild type (WT) hearts and 655 genes were downregulated. Reactome, KEGG, and Gene Ontology (GO) pathway enrichment performed with Enrichr.
g) Venn diagram of differentially expressed genes (DEGs) in female *Chchd10*  versus WT mice. Bulk RNA-seq was performed on cardiac biopsies (n=3) at 14 weeks of age. 681 genes were upregulated in female *Chchd10* vs wild type (WT) hearts and 567 genes were downregulated. Reactome, KEGG, and Gene Ontology (GO) pathway enrichment performed with Enrichr.
h) Quantification of mitochondrial DNA (mtDNA) in cardiac biopsies from male WT (blue, n=3), *Chchd10* (red, n=3) mice at 7, 14, and 35 weeks. Primers directed at mtDNA encoding 16s rRNA and β-actin for nDNA were used. Data are means ± SEM, multiple unpaired t-test P values indicted.
i) Whole-body deletion of *Sting* in *Chchd10* mice. Generation of the *Chchd10^S55L/+^Sting^Gt/Gt^* mice were generated by intercrossing *Chchd10^S55L/+^* (*Chchd10*) with *Sting^Gt/Gt^* mice lacking functional STING (*Sting*). Created with Biorender.com
j) Left ventricular ejection fraction (% LVEF) of *Sting* (*Sting^Gt/Gt^*, orange, n=6) and *Chchd10/Sting* (*Chchd10^S55L/+^Sting^Gt/Gt^*, purple, n=8) male mice at 38 weeks (left) and 48 weeks (right) of age. Data represent mean *±* SEM. Student’s t-test, **P<0.01, ns=not significant. Dotted line represents *Chchd10* %LVEF at 35 weeks.
k) Body mass of *Sting* (*Sting^Gt/Gt^*, orange, n=3-12), *Chchd10/Sting* (*Chchd10^S55L/+^Sting^Gt/Gt^*, purple= n=3-14) male mice. Data are means ± SEM, 2-tailed unpaired Student’s t test used to identify significant differences (P<0.05). Dotted grey line represents the age after which body mass differences between WT and *Chchd10* male (17 weeks) mice.
l) Kaplan-Meier survival curve of *Sting* (orange, n=14), *Chchd10/Sting* (purple= n=10) male mice. Dotted line (grey) represents median lifespan of *Chchd10* mice Log-rank test *Chchd10* vs *Chchd10/Sting*, P=0.002.

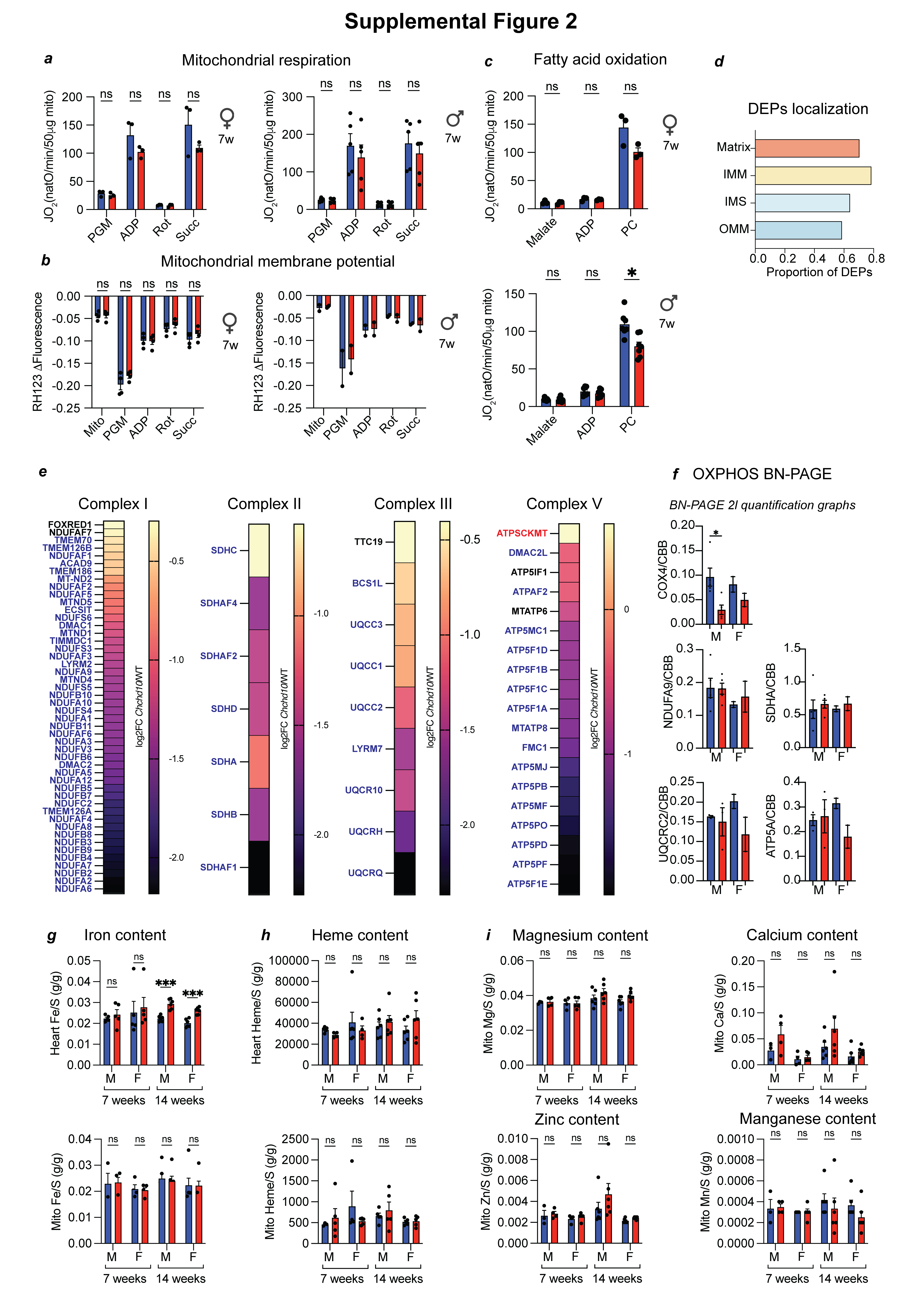

**Supplemental Figure 2 – Impaired mitochondrial respiration in mutant *Chchd10* hearts**

a) Oxygen consumption rates (JO_2_) of cardiac mitochondria isolated from WT (n=3-5) and *Chchd10* (n=3-5) female (left) and male (right) mice at 7 weeks. JO_2_ measured sequentially in the presence of pyruvate, glutamate, malate (PGM), adenosine diphosphate (ADP), rotenone (Rot), and succinate (Succ). Data represent mean ± SEM; multiple unpaired t-test, ns=not significant.
b) Mitochondrial membrane potential (ΔΨ) measured by quenching of Rhodamine 123 (RH123) fluorescence in cardiac mitochondria of WT and *Chchd10* male (n=5) and female (n=2) mice from Figure S2a. Data represent mean ± SEM; multiple unpaired t-test, ns=not significant.
c) Oxygen consumption rates (JO_2_) of cardiac mitochondria isolated from WT (n=3-7) and *Chchd10* (n=3-7) female (top) and male (bottom) mice at 7 weeks. JO_2_ measured sequentially in the presence of malate, adenosine diphosphate (ADP), and palmitoyl carnitine (PC). Data represent mean ± SEM; multiple unpaired t-test, *P<0.05, ns=not significant.
d) Mitochondrial localization of DEPs in Figure 2g according to MitoCarta 3.0 represented as proportion of total quantified DEPs in each mitochondrion subcompartment.
e) Heatmap of Complex I, II, III, and V proteins quantified by proteomics in Figure 2g and significantly upregulated (red), downregulated (blue) or unchanged (black) between WT (n=5) and *Chchd10* (n=5) male mice at 14 weeks (Supplemental Dataset 5).
f) Densitometric quantification of OXPHOS complexes in Figure 2l is relative to Coomassie brilliant blue (CBB). Data are means ± SEM, 2-tailed unpaired Student’s t test. * P<0.05.
g) Iron content in total heart (top, n=5-6) and cardiac mitochondria (bottom, n=3-6) samples measured by inductively coupled plasma–optical emission spectrometry (ICP-OES) from WT and *Chchd10* male (M) and female (F) mice at 7 and 14 weeks of age. Data represent mean values normalized to sulfur (S) *±* SEM, multiple unpaired t-test, *** P<0.001, ns=not significant.
h) Heme content in total heart (top, n=5-6) and cardiac mitochondria (bottom, n=3-6) samples measured by HPLC from WT and *Chchd10* male (M) and female (F) mice at 7 and 14 weeks of age. Data represent mean values normalized to sulfur (S) *±* SEM, multiple unpaired t-test, ns=not significant.
i) Magnesium, Zinc, Calcium, and Manganese content in cardiac mitochondria measured by inductively coupled plasma–optical emission spectrometry (ICP-OES) from WT and *Chchd10* male (M) and female (F) mice at 7 and 14 weeks of age. Data represent mean values normalized to sulfur (S) *±* SEM, multiple unpaired t-test, ns=not significant.

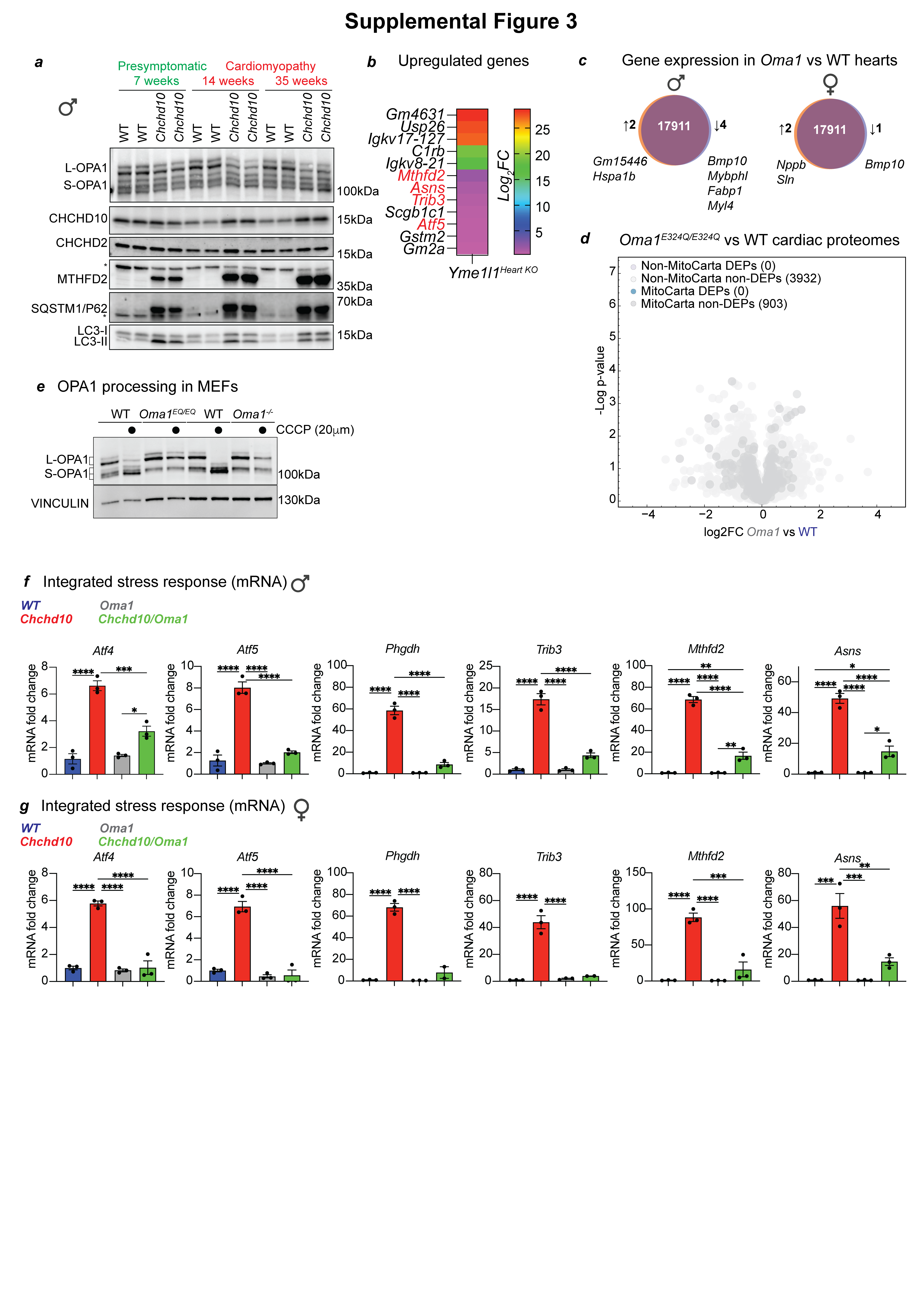

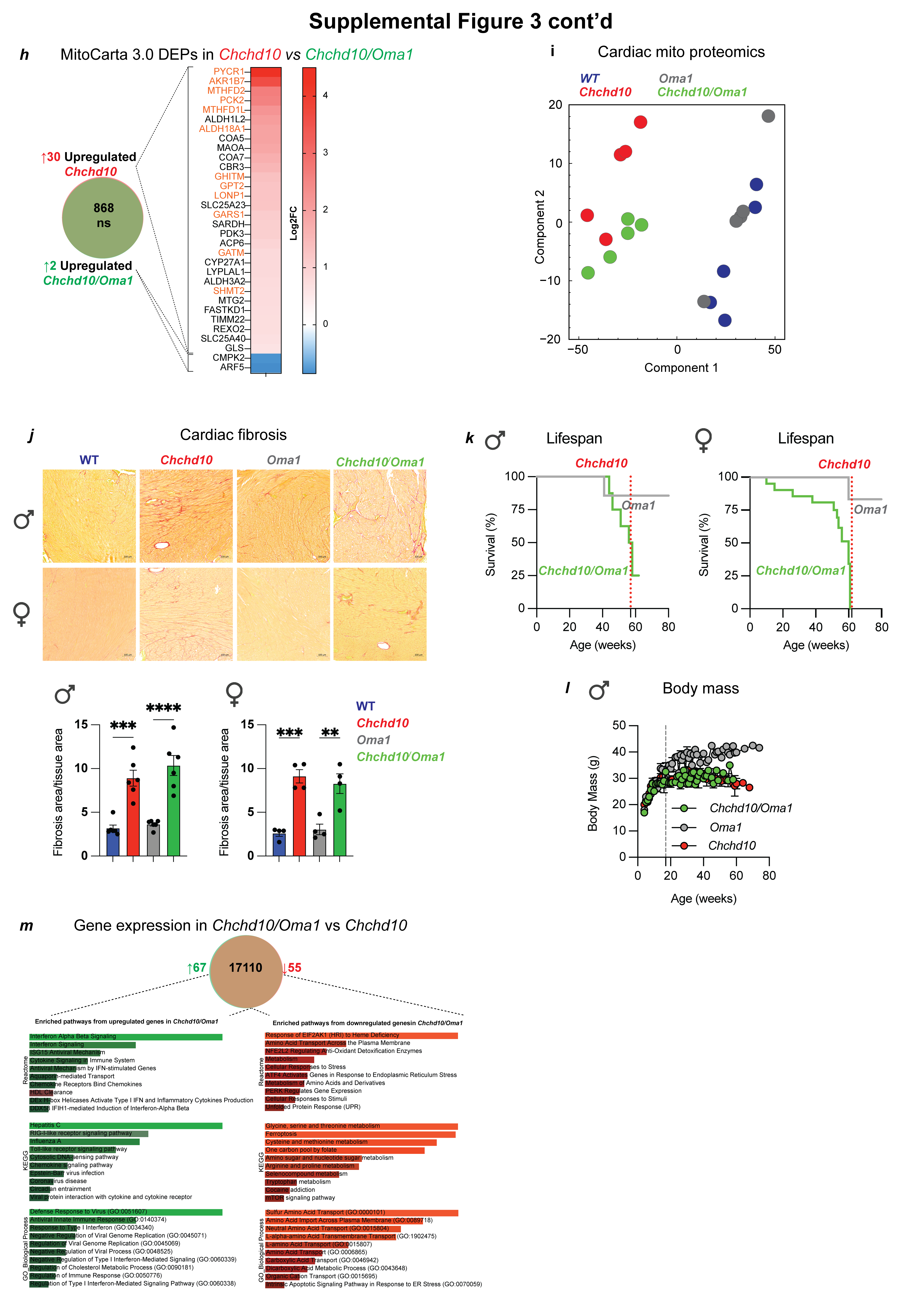
**Supplemental Figure 3 –OMA1 and mtISR activation**

a) Immunoblots in cardiac lysates of presymptomatic (green) and symptomatic (red) WT and *Chchd10* mice.
b) Heatmap of upregulated differentially expressed genes (DEGs) in cardiomyocyte-specific *Yme1l* knockout mice (*Yme1l1^HeartKO^*). Bulk RNA-seq was performed on cardiac biopsies from male (n=3) mice compared to sex-matched littermate controls at 35 weeks of age. Integrated stress response (ISR) genes are highlighted red.
c) Venn diagram of differentially expressed genes (DEGs) in *Oma1* mice. Bulk RNA-seq was performed on cardiac biopsies from male (n=3) and female (n=3) *Oma1*  mice compared to sex-matched WT controls at 14 weeks of age. DEG is defined by Log2FC>2 and padj <0.01 (Supplemental Dataset 1).
d) Volcano plot of cardiac proteomics WT vs Oma1 No differentially expressed proteins (DEPs) were identified. Significance is considered a permutation based-FDR at 0.05 highlighted by color.
e) Immunoblots of OPA1 processing in mouse embryonic fibroblasts (MEFs) derived from wild type (WT) and *Oma1^E324Q/E324Q^* (*Oma1^EQ/EQ^*) mice. Carbonyl cyanide m-chlorophenyl hydrazine (CCCP) used to induce stress-induced L-OPA1 processing. WT and OMA1 knockout (*Oma1^-/-^*) MEFs were used as a control^59^.
f) Analysis of integrated stress response (ISR) genes via qRT-PCR of total cardiac biopsies from wild type (WT, blue, n=3), *Chchd10* (red, n=3), *Oma1* (grey, n=3), and *Chchd10/Oma1* (green, n=3) male mice at 14 weeks in Figure 3c. Data are relative mean fold changes ± SEM, 2-tailed unpaired Student’s t test. *P<0.05, **P<0.01, ***P<0.001,****P<0.0001.
g) Analysis of integrated stress response (ISR) genes via qRT-PCR of total cardiac biopsies from wild type (WT, blue, n=3), *Chchd10* (red, n=3), *Oma1* (grey, n=3), and *Chchd10/Oma1* (green, n=2-3) female mice at 14 weeks in Figure 3c. Data are relative mean fold changes ± SEM, 2-tailed unpaired Student’s t test. *P<0.05, **P<0.01, ***P<0.001,****P<0.0001.
h) Heat map of differentially expressed proteins (DEPs) upregulated in cardiac mitochondria profiled from *Chchd10* (red, n=5) versus *Chchd10/Oma1* (green, n=5) hearts by mass spectrometry (Supplemental Dataset 5).
i) Principal component analysis (PCA) of cardiac proteomics performed on wild type (WT, n=5, blue), *Chchd10* (n=5, red), *Oma1* (n=5, grey) and *Chchd10/Oma1* (n=5, green) male mice at 14 weeks of age (Supplemental Dataset 5).
j) Cardiac histology of 4 genotypes and both sexes. Representative sirius red myocardium staining of wild type (WT), *Chchd10*, *Oma1*, and *Chchd10/Oma1* male (22 weeks of age, n=6) and female (22 weeks of age, n=4) mice. Tissue fibrosis analysis was performed using an automated macro developed in QuPath and quantified (bottom). Data represent mean *±* SEM. One-way ANOVA, * P<0.05, **P<0.01, ****P<0.0001, ns=not significant.
k) Kaplan-Meier survival curve (left) of *Oma1* (grey, n=15), and *Chchd10/Oma1* (green, n=15) male mice and (right) *Oma1* (grey, n=13), and *Chchd10/Oma1* (green, n=21) female mice. Dotted red line represents median lifespan of *Chchd10* mice. Log-rank test *Chchd10* vs *Chchd10/Oma1*; male P=0.0844 and female P=0.2520.
l) Body mass of *Oma1* (grey), and *Chchd10/Oma1* (green) male mice compared to data from male *Chchd10* mice. Dotted grey line represents the age after which body mass differences between WT and *Chchd10* male (17 weeks) mice are observed.
m) Venn diagram of differentially expressed genes (DEGs) in female *Chchd10/Oma1*  versus *Chchd10* mice. Bulk RNA-seq was performed on cardiac biopsies (n=3) at 14 weeks of age. Reactome, KEGG, and Gene Ontology (GO) pathway enrichment performed with Enrichr.

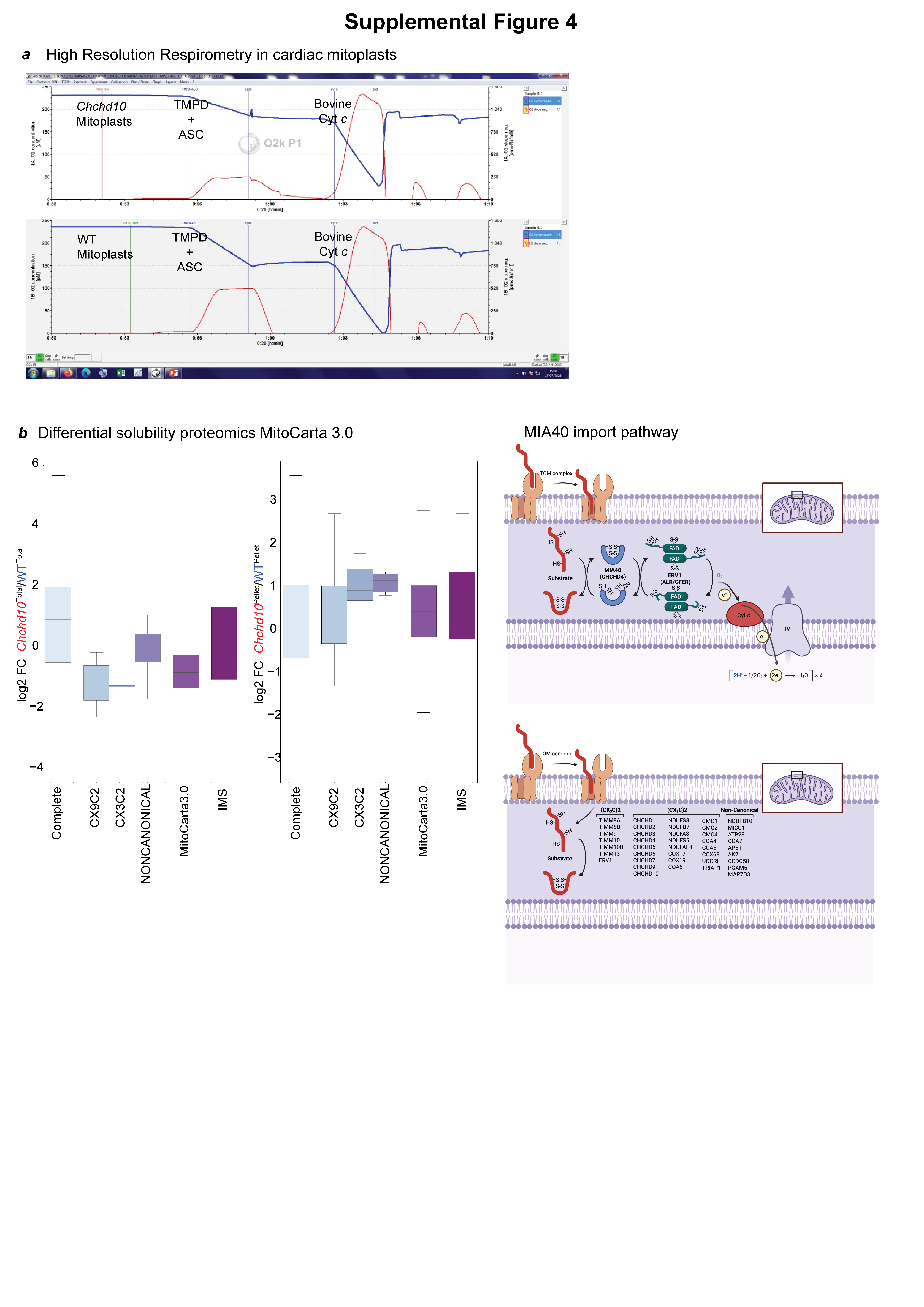
**Supplemental Figure 4 – Differential solubility proteomics in *Chchd10* mice**

a) Representative traces of oxygen consumption rates (JO_2_) measured in *Chchd10* and wild type (WT) mitoplasts using High Resolution Respirometry (Oroboros). Blue trace represents chamber oxygen (O_2_) concentration (μM, left y-axis) and red trace represents O_2_ consumption (pmol/(s*ml, right y-axis). JO_2_ measured sequentially following addition of N,N,N′,N′-Tetramethyl-p-phenylenediamine (TMPD) plus ascorbate (ASC) and bovine cytochrome *c* (Cyt *c*) at indicated times (running time in hours: min (h:min), x axis).
b) Boxplots of MitoCarta, IMS, and MIA40 client types based on differential solubility proteomic analyses of total (T, left) and pellet (P, right) fractions of detergent-solubilized cardiac mitochondria from WT and *Chchd10* male mice at 14 weeks of age (left, Supplemental Dataset 7). The whiskers show the minimal and maximal values. The box edges indicate the first (25%) and third (75%) quantile. The black line in the box indicates the median (50%).Cartoon representation of MIA40/ERV1 import pathway (top) in which client proteins^2^ are indicated (right). Created with Biorender.com
